# Thermosensory pathway makes a U-turn to respond to heat

**DOI:** 10.64898/2026.11.06.764838

**Authors:** Luciana Bianchimano, Cha San Koh, María Belén De Luca, Germán Murcia, Matías Ezequiel Pereyra, Anna B. Hamacher, Matias D. Zurbriggen, Jorge J. Casal

**Author notes:** Author for correspondence: Jorge J. Casal. These authors contributed equally to this work (Cha San Koh, María Belén De Luca, Germán Murcia). L’Institut des Sciences des Plantes, Universidad Paris-Saclay. 630 Rue Noetzlin Bâtiment, 91190 Gif-sur-Yvette, Paris, France (María Belén De Luca).

## Abstract

Plant organs exhibit a unimodal growth response to temperature, yet the mechanistic links connecting growth-promoting and growth-inhibiting thermal ranges remain unresolved. In *Arabidopsis* seedlings, warm temperatures promote hypocotyl elongation by relieving phytochrome B (phyB) inhibition, thereby activating the transcription factors PIF4 and PIF7 to drive auxin synthesis. Here, we demonstrate that temperatures beyond the growth optimum actively suppress growth by downregulating auxin levels, auxin signalling, and PIF4 abundance specifically within cotyledons, the primary site of auxin production. Strikingly, whereas warm temperatures induce the shrinking of phyB nuclear bodies, heat stress causes phyB to assemble into enlarged condensates, even when phyB is predominantly in its inactive form. We recapitulated heat-induced phyB condensates in mammalian cells, demonstrating an intrinsic biophysical mechanism. Together, our findings indicate that heat-induced growth inhibition is not merely passive stress damage, but an active molecular reversal of the thermomorphogenic promotion pathway mediated by distinct phyB condensates.

## Introduction

Growth of plant organs typically follows a unimodal curve, increasing with temperature to an optimum and declining at higher temperatures (Parent & Tardieu, 2012; Zhu *et al*., 2022). The optimum growth temperature is generally higher for species originating from tropical regions than for those from temperate regions (Parent & Tardieu, 2012). Even within a species, organs differ in their thermal responses. In *Arabidopsis thaliana*, hypocotyl elongation increases between 10°C and 30°C (Legris *et al*., 2017), primary root elongation increases between 10°C and 25°C (Ibañez *et al*., 2017; Yang *et al*., 2017; Borniego *et al*., 2022) whereas cotyledon expansion can decline between 20°C and 28°C (Hahm *et al*., 2020; Costigliolo Rojas *et al*., 2022).

Our understanding of the mechanisms involved in the promotion of hypocotyl growth by warm temperatures has improved substantially during the last decades (Quint *et al*., 2023). Warm temperatures enhance auxin synthesis in the cotyledons, followed by hormone transport to the hypocotyl to drive elongation (Gray *et al*., 1998; Franklin *et al*., 2011; Bellstaedt *et al*., 2019). The bHLH transcription factors PHYTOCHROME INTERACTING FACTOR 4 (PIF4) (Franklin *et al*., 2011; Sun *et al*., 2012) and PIF7 (Fiorucci *et al*., 2020) activate auxin biosynthesis genes. PIF7 accumulates via warm temperature-dependent translational regulation (Fiorucci *et al*., 2020; Chung *et al*., 2020), while *PIF4* increases via transcriptional (Koini *et al*., 2009) and post-translational (Foreman *et al*., 2011; Zhang *et al*., 2025; Tan *et al*., 2026) controls. Upstream, the EVENING COMPLEX (EC) represses *PIF4* expression at night (Nusinow *et al*., 2011), but one of its components, EARLY FLOWERING 3 (ELF3), undergoes temperature-induced phase separation into nuclear condensates that prevent the EC formation (Jung *et al*., 2020; Silva *et al*., 2020). PIF5 and PIF3 further contribute to thermomorphogenesis (Romero-Montepaone *et al*., 2021).

The photoreceptor phytochrome B (phyB) represses PIF4 accumulation and intrinsic ability to bind to transcriptional targets (Park *et al*., 2018; Pham *et al*., 2018). Warm temperatures accelerate thermal reversion of phyB from the biologically active far-red light (FR)-absorbing Pfr form to its inactive red light (R) absorbing Pr form (Jung *et al*., 2016; Legris *et al*., 2016). phyB Pfr assembles into nuclear condensates or nuclear bodies (NBs) through liquid-liquid-phase separation (Kircher *et al*., 1999; Yamaguchi *et al*., 1999; Chen *et al*., 2003, 2022). These NBs dissociate upon FR irradiation (Rausenberger *et al*., 2010; Van Buskirk *et al*., 2014) and shrink at elevated temperatures (Legris *et al*., 2016; Hahm *et al*., 2020; Chen *et al*., 2022). phyB NBs are multi-protein condensates containing both primary and secondary interacting partners (Kim *et al*., 2023). Although phyB is able to form condensates when expressed in mammalian cells in the absence of other plant proteins (Chen *et al*., 2022), the size of these condensates increases substantially in the presence of interacting partners(Beyer *et al*., 2015b; Chen *et al*., 2022). In plant cells, NBs formation also depends on several such factors, including PHOTOPERIODIC CONTROL OF HYPOCOTYL 1 (PCH1) and PCH1-LIKE (PCHL) (Galvão *et al*.; Enderle *et al*., 2017; Huang *et al*., 2019; Fang *et al*., 2022).

Although warm temperatures promote hypocotyl growth through well-understood mechanisms, how heat represses growth remains elusive. This inhibition could reflect cellular damage (such as membrane dysfunction, protein denaturation, or reactive oxygen species) or an actively regulated pathway. One hypothesis is that heat further amplifies warm-temperature growth pathways, causing auxin-induced apoplastic acidification to reach supra-optimal intensities known to inhibit hypocotyl growth (Wang *et al*., 2025). Instead, we find that heat reverses key processes underlying hypocotyl thermomorphogenesis. Because warm temperatures accelerate phyB thermal reversion and promote NBs disassembly, phyB condensates would be expected to decline at higher temperatures. Unexpectedly, quantitative imaging in plants, together with an orthogonal reconstitution system in mammalian cells, reveals that heat enlarges phyB condensates, uncovering a temperature-dependent reorganization of phyB associated with growth repression.

## Materials and methods

### Plant material

We used *Arabidopsis thaliana* seedlings of the genotypes listed in Table S1. Seeds were sown on 1 % agar-water in clear plastic boxes of 3 cm length, 4 cm width and 2 cm height, and stratified 4 d at 5 °C in darkness. Then, the boxes were transferred to a growth room at 22°C with a photoperiod of 10 h of 80-90 μmolm^−2^ s^−1^ of white light (400–700 nm) provided by a mixture of fluorescent and halogen lamps (red/far-red ratio = 1.1). In hypocotyl-growth experiments, the seedlings were grown parallel to the agar surface, which was placed vertically. For other determinations, the seedlings were grown with their hypocotyls perpendicular to the agar surface, which was placed horizontally. For luminometer experiments, the seedlings were grown in 96-well plates containing 1 % agar-water. One hour after the beginning of the fourth day (time= 0 h), the seedlings were transferred to the different treatments.

### Temperature treatments

Unless indicated otherwise, 1 h after the beginning of the fourth day (time= 0 h), the seedlings were transferred to the different temperatures (10°C, 15°C, 22°C, 28°C, 33°C or 37°C), where they remained 9 h (hypocotyl growth), 4h or 8h (other observations) without changes in light conditions.

In selected experiments, the seedlings were transferred at time= 0 h to different combinations of temperature (28°C, 33°C or 37°C) and R pulses (5 min, 35 μmol m^-2^ s^-1^) or FR pulses (5 min, 15 μmol m^-2^ s^-1^) followed by darkness, hourly R pulses or to continuous high-irradiance R (175 μmol m^-2^ s^-1^). The R and FR treatments (wavelengths of maximum emission at 660 nm and 730 nm, respectively) were provided by LEDs. The specific sequence of conditions is detailed along with the description of the results.

### Rate of hypocotyl elongation

For each biological replicate, 10, 12 or 6 seedlings (see figure legends) were photographed from the side with a digital camera (PowerShot; Canon, Tokyo, Japan) at the beginning of treatments (time= 0 h) and at the end of the photoperiod (time= 9 h) of the fourth day. Hypocotyl length was determined at both time points using image-processing software (Sellaro *et al*., 2009) and used to calculate the increments over the 9-h period.

### Quantifying genetic contributions to hypocotyl growth

Hypocotyl growth (G) was modeled as the ratio of maximum growth capacity (G_max_) to the total inhibitory activity (I) plus a constant of one, representing growth in the absence of inhibition:

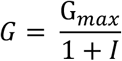

In this model, total inhibitory activity in the wild type is the sum of the contribution from gene X (I_X_) and other systemic components (I_other_). In the gene *X* mutant, inhibitory activity is attributed solely to these other components. Therefore, I_other_ was first calculated using the growth equation for the mutant:

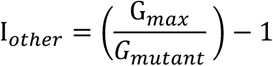

This value was then substituted into the wild-type equation to isolate the specific inhibitory contribution of gene *X*:

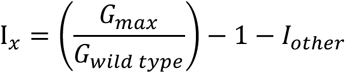

Because *pifs* and *bes1/bzr1* mutants exhibit reduced growth relative to the wild type, their calculated inhibitory contributions yielded negative values. To reflect their biological roles as growth enhancers, these effects are presented as promotion, yielding positive values. Maximum growth capacity (G_max_) was defined using *phyB* mutant data from plants grown under shade at 28°C.

### Confocal microscopy

Confocal images from Arabidopsis seedlings were taken from live samples except in the case of the experiments where light pulses were followed by darkness to analyse phyB-GFP NBs. In the latter cases, to avoid any spurious exposure to light that could distort the results, the seedlings were fixed by immersion in 1% paraformaldehyde in 1× PBS, applying a vacuum for 24 seconds, followed by two 30-minute washes with NH_4_Cl and suspension in 1× PBS, as described (Pereyra *et al*., 2023). In all cases, images were taken of the epidermis or the first sub-epidermal layers, taken 4 h and 8 h after the application of treatments of temperature. The organs observed (cotyledons or hypocotyls) are indicated in each case.

For the analysis of the *AuxSen* (Herud-Sikimić *et al*., 2021) and Apo-pHusion (Gjetting *et al*., 2012) reporters, confocal fluorescence images were taken with the LSM 510 Meta (Zeiss) laser scanning confocal microscope with an Plan-Apochromat 40×/1.2 objective lens. For FRET ratio measurement of *AuxSen*, the donor fluorophore was excited with a 405 nm wavelength laser bean, and the emission of the donor was detected between 460 nm and 500 nm, while the emission of the acceptor (FRET) was detected between 505 nm and 550 nm. Additionally, to verify its performance, the acceptor was excited with a 488 nm wavelength laser, and its emission was detected between 505 nm and 550 nm. Data correspond to the ratio between the emission intensities of the acceptor (505-550 nm) and donor (460-500 nm). For Apo-pHusion, we used a sequential scanning of EGFP and mRFP1 proteins. EGFP was excited at 488 nm and detected between 500 nm and 550 nm while mRFP1 was excited at 585 nm and detected between 600 nm and 630 nm, as described (Gjetting *et al*., 2012). Data correspond to the ratio between EGFP and mRFP1 fluorescence intensities.

To quantify nuclear fluorescence, a Zeiss LSM 710 NLO confocal microscope was used, which allows the separation of the signal emitted by fluorescent proteins from that of chloroplasts. For visualisation of proteins fused to GFP and YFP, mCitrine, and DII-VENUS, we used an Argon excitation laser (λ = 488 nm), and fluorescence was detected using a BP 505-530, 505-550, and 505-570 filter, respectively. Since chloroplasts exhibit strong autofluorescence, and chlorophylls are excited and fluoresce in wavelength ranges within those used for the excitation and detection of fluorescent proteins, a separate channel with a He-Ne excitation laser (λ = 543 nm) and an LP 630 filter was used to detect chlorophyll and distinguish chloroplasts in the fluorescent protein channel.

Nuclear fluorescence is expressed per cell as the ratio between the integral of nuclear fluorescence and the number of cells within the area under analysis. Their spindle shape, compressed against the cell wall, identified nuclei and in many cases, the presence of the nucleolus was detected. The average fluorescence intensity within the nuclei and the number of nuclei within the area under analysis were recorded by using Fiji software (Schindelin *et al*., 2012), and their product defined the integral of nuclear fluorescence within the area under analysis. The area per cell was measured in 5-10 cells and the total area under analysis were measured with the aid of the transmission and chlorophyll channels, and these data were used to calculate the number of cells within the area under analysis.

To analyse phyB NBs, we used LSM 510 Meta (Zeiss) to first obtained the image of the area to be analysed, and then performed 8× magnifications of that image in the areas of interest focused on one nucleus to optimise the visualisation of NBs. The area, number and fluorescence of the NBs were obtained by using an automatic analysis tool developed in Icy (de Chaumont *et al*., 2012; Legris *et al*., 2016) by using Fiji software (Schindelin *et al*., 2012).

### Quantitative RT-PCR

For each biological replicate, approximately 50 mg of Col-0 seedlings were harvested, frozen in liquid nitrogen, and ground with a mortar and pestle to a fine powder, 4 h after the beginning of the temperature treatments. Total RNA was extracted using Trizol reagent (Invitrogen) according to the manufacturer’s instructions. One μg of RNA was treated with RQ1 RNase-Free DNase (Promega) and reverse transcribed using Superscript II Reverse Transcriptase (SSII RT) (Thermo Fisher Scientific) and oligo-dT, according to the manufacturer’s protocol. The resulting cDNAs were then amplified using an Mx3000P Real Time Cycler PCR System (Agilent Technologies). The *UBQ2* gene transcript (AT2G36170) was used as a reference. Quantitative RT-PCR analysis was performed using the standard curve method (Agilent Technologies, 2012). Primers are listed in Table S1.

### Colorimetric auxin quantification

Auxin levels were quantified following a colorimetric method (Manna *et al*., 2024). For each biological replicate, 50-100 mg fresh weight of Col-0 seedlings were harvested 4 h after the onset of temperature treatments, weighed, and flash-frozen in liquid nitrogen. Samples were ground to a fine powder using a mortar and pestle and extracted in 1 mL of absolute ethanol. The mixture was swirled to ensure IAA solubilisation and incubated on ice for 10 min, followed by centrifugation at 1,000× *g* for 10 min at 4 °C. For the colorimetric reaction, 80 µL of the resulting supernatant was combined with 100 µL of Salkowski reagent (distilled H_2_O, concentrated H_2_SO_4_, and 0.5 M FeCl_3_ in a 25:15:0.75 ratio) in a 96-well transparent microplate. Following a 30-min incubation at room temperature in the dark, absorbance was measured at 540 nm using a Varioskan LUX multimode reader (Thermo Fisher Scientific). IAA concentrations were determined using a standard calibration curve (range: 0.49–501.76 mg/L, Table S1) and normalised to the fresh weight of each sample.

### Luminometer readings

Transgenic seedlings bearing the LUC reporter were incubated with 30 μL of 0.5 mM D-luciferin per well 24 h before starting to record luciferase activity as described (Murcia *et al*., 2022). Luciferase (LUC) activity was recorded with a Varioskan™ LUX multimode microplate reader (Thermo Fisher Scientific) for three seconds, 4 h after the beginning of the temperature treatments.

### Protein blots

For each biological replicate, approximately 50 mg of seedlings were harvested and frozen in liquid nitrogen and ground with a mortar and pestle to a fine powder, 4 h or 8 h after the beginning of the temperature treatments. Blot reactives are listed in Table S1. Total protein was extracted on a buffer containing 125 mM Tris-HCl pH 7.4, 0.2% SDS, 10% glycerol, 6 M urea, and 1% β-mercaptoethanol(Nieto *et al*., 2015). Samples were heated for 3 min at 95°C, then centrifuged for 15 min at 13,000 rpm, and the supernatant was removed. Proteins were separated by PAGE using 12% (phyB, PIF7) or 8% (PIF4) gels. Gels were then electrotransferred onto a 0.45 µm nitrocellulose membrane (Thermo Fisher catalogue 88018) using a wet transfer system for 2 h at 100 V. Rabbit polyclonal anti-HA and mouse polyclonal anti-GFP (Thermo Fisher catalogue PA1985 and A-11122) were used as primary antibodies. Anti-rabbit IgG (H+L) with HRP (Thermo Fisher catalogue 32460) and anti-mouse (H+L) were used as secondary antibodies. Detection was carried out using the SuperSignal^TM^ West Pico PLUS Chemiluminescent Substrate (Thermo Fisher Scientific catalogue 34580) and X-ray films. Ponceau red staining was used as a control for loading uniformity. Band quantification was carried out using the Fiji program, always selecting the same rectangle size. Full blots corresponding to each biological replicate are shown in Figures S1, S2 and S3. The intensity of each protein band was divided by the intensity of the Ponceau band to calculate its relative abundance (see Dataset 1 for the data corresponding to the quantification and normalisation of each band).

### β-Glucuronidase (GUS) activity measurements

For each biological replicate, approximately 20 transgenic seedlings carrying the *pDR5:GUS* reporter were harvested 4 h after the start of temperature treatments. Samples were ground at ambient temperature in 200 µL of extraction buffer (250 mM phosphate buffer pH 7, 0.5 M Na_2_EDTA, pH 8, 10% SDS, 10% Triton X-100, and 14.4 M β-mercaptoethanol) and immediately placed on ice. Extracts were clarified by centrifugation at 12,000 rpm for 5 min (4 °C). To measure activity, 5 µL of supernatant was combined with 195 µL of reaction buffer (extraction buffer supplemented with 25 mM 4-methylumbelliferyl β-D-glucuronide [4-MUG]) in a 96-well microtiter plate. The plates were incubated at 37 °C for 1.5 h in darkness before the reaction was terminated using a carbonate stop buffer (Na_2_CO_3_ and NaOH). Fluorescence was determined at an excitation wavelength of 365 nm and an emission wavelength of 455 nm using a Varioskan LUX reader (Thermo Fisher Scientific) (Blázquez, 2007). Activity was quantified using a standard curve of 4-methyl umbelliferone (4-MU) and normalised to the number of seedlings.

### Conventional confocal microscopy of mammalian cells

Chinese hamster ovary cells (CHO-K1) were co-transfected with the *pSV40:PHYB-eGFP* and *pSV40:PIF3-mCerulean, pSV40:PIF4-mCerulean* or *pSV40:PIF7-mCerulean* plasmids (cell and recombinant materials are listed in Table S1). The plasmids were constructed using Aqua cloning(Beyer *et al*., 2015a) and documented using GMOCU (Wagner *et al*., 2024). CHO-K1 cells were cultured in Ham’s F12 medium supplemented with 10% (v/v) fetal bovine serum (FBS) and 1% (v/v) penicillin/streptomycin. Twenty-four hours prior to transfection, 50,000 cells per well were seeded onto coverslips in 24-well plates in 500 μl of medium. For transfection, 0.75 μg of DNA was diluted in 50 μl of Opti-MEM and combined with a PEI/Opti-MEM mixture [2.5 μl of PEI (1 mg/ml) in 50 μl of Opti-MEM per well] (Blomeier *et al*., 2021). Following a brief incubation, 100 μl of the transfection complex was added dropwise to each well. The medium was replaced after 4 h to minimise cytotoxicity. Twenty-four hours post-transfection, cells were incubated with 15 μM PCB chromophore (Beyer *et al*., 2015b) for 1 h before applying the light and temperature treatments specified in the figure legends.

The cells were fixed in the dark using 4% PFA (10 min on ice, followed by 10 min at room temperature), with PBS washes performed before and after fixation. Nuclei were stained with 0.1% DAPI for 5 min at room temperature. Finally, coverslips were mounted onto microscope slides using 8 μl of Mowiol/DABCO. The images were analysed as described for plant samples.

### High-content imaging of transfected mammalian cells

The three constructs used for high-content imaging experiments using CHO-K1 cell line are *pSV40::PIF3-eGFP::tSV40* (pMZ725)^33^, *pSV40::PHYB-mCherry::tSV40* (pSJ025)^33^ and *pSV40::PHYB^Y276H^-mCherry::tSV40* (pCSK208A).

CHO-K1 cells were initially maintained in Ham’s F12 medium (PAN Biotech, Aidenbach, Germany; P04–14500) supplemented with 10% (v/v) tetracycline-free FBS and 1% (v/v) penicillin/streptomycin (PAN Biotech). They were then gradually adapted over 2 weeks to 75% CO₂-independent medium (Gibco #18045054) supplemented with GlutaMAX™ (Gibco #35050061), 10% FBS, 1% penicillin/streptomycin, tetracycline and 25% Ham’s F12 to support CHO-K1 growth. This CO₂-independent formulation, lacking HEPES, maintains physiological pH without external CO₂. Experiments were performed at 15°C, 22°C, 28°C, and 37°C. Adapted CHO-K1 cells (about 14,000 cells in 150 μl medium per well) were seeded into PhenoPlate™ 96-well microplates (Revvity) and incubated for 1 day at 37°C with 5% CO₂. For transfection in 96-well format, 0.2 μg total DNA per well was diluted in Opti-MEM™ with P3000™ reagent (2 μl μg^-1^ DNA) to a final volume of 5 μl. The diluted DNA was mixed 1:1 with diluted Lipofectamine™ 3000 in Opti-MEM™ to form a 10 μl DNA– lipid complex, incubated for 10–15 min at room temperature, and added to cells. Transfected cells were incubated for 24 h at 37°C with 5% CO₂. Although removal of transfection complexes is not required, a medium exchange 4–5 h post-transfection improved imaging quality by reducing background. The following day, 15 μM phycocyanobilin (PCB; Frontier Scientific, P14137-50mg) was added and cells were incubated for 1 h at 37°C in darkness. Plates were then illuminated with red light (660 nm peak, 20 μmol·m⁻²·s⁻¹) for 15 min at the indicated temperatures (15°C, 22°C, 28°C, 37°C) in temperature-controlled chambers. Under continuous red light, the medium was rapidly exchanged with chromophore-free medium pre-equilibrated to the respective temperature. Cells were then incubated in complete darkness at the same temperature for 60 min. All subsequent steps were performed under dim green light to mimic dark conditions. Cells were washed once with 100 μl pre-equilibrated PBS, then fixed in 100 μl 4% paraformaldehyde (PFA, Roth) for 15 min on ice, protected from light. After removing PFA, cells were washed once with 100 μl PBS at room temperature. To enable multiplexing in high-content analysis, two nuclear stains were used. Cells were first incubated with 50 μl HCS NuclearMask™ Deep Red (Thermo Fisher H10294) for 45 min at room temperature, protected from light. After one PBS wash, nuclei were counterstained with 50 μl DAPI (2 μg/ml) for 15 min. After removal of DAPI, cells were overlaid with 100 μl PBS and stored at 4°C until imaging.

To ensure quantitatively unbiased and reproducible observations, a fully automated high-content imaging workflow was implemented for fixed CHO-K1 cells. Imaging was performed on an Operetta CLS High Content Imaging System (Revvity) at the Center for Advanced Imaging, Heinrich Heine University Düsseldorf, equipped with a 4.7 MP sCMOS sensor (resolution of 2160 × 2160 pixels). Excitation/emission settings were configured as follows: DAPI (355–385 nm / 430–500 nm; Nuclear Counterstain), eGFP (460–490 nm / 500–550 nm; PIF3 condensates), mCherry (530–560 nm / 570– 650 nm; phyB condensates), and HCS NuclearMask Deep Red (615–645 nm / 665–760 nm).

Data acquisition was accelerated using the PreciScan module in Harmony™. An initial low-magnification pre-scan (20× air objective, NA 0.4, binning 3) was used to identify cells containing condensates, followed by automated image analysis using an optimised Harmony™ pipeline (Fig. S4). High resolution imaging at 40x was performed using a custom Harmony™ analysis sequence that integrated automated cell selection, stage repositioning, autofocus, and confocal z-stack acquisition based on candidates identified during the pre-scan.

Confocal z-stacks were acquired using a 40× air objective (NA 0.75, binning 1), centred on the identified cells. Per well, 100 – 200 fields were imaged depending on transfection efficiency. Z-stacks comprised five optical sections for cells transfected with two plasmids and four optical sections for cells transfected with a single plasmid, covering a z-range of 1.2-4.2 μm across the evaluated cells. Non-transfected cells, defined by reporter fluorescence below a predefined threshold, were automatically excluded. Nuclear and cytoplasmic segmentation were performed using NuclearMask Deep Red and DAPI signals, respectively, while condensate detection was based on the fluorescence channels described above.

Images were processed as maximum intensity projection (MIP) for condensate quantification using a custom pipeline in Harmony™ version 5.2. The complete customised sequence analysis pipelines are available from the authors upon reasonable request.

### KNIME-based data processing workflow

Data analysis was performed using a customized workflow (Fig. S4) in KNIME Analytics Platform (v5.11.0; www.knime.com), integrating Harmony™ image analysis outputs for segmentation and feature extraction. Condensates were deconvoluted based on fluorescence properties. Cells in contact with the image borders were excluded from the image analysis. Quality control was performed in KNIME by applying cell-size thresholds to exclude artefacts and debris, as well as filtering based on condensate signal intensity relative to the background. Condensates were subsequently classified by subcellular localization using features derived from DAPI and NuclearMask staining. A metadata harmonization step followed, whereby cell-level information was systematically assigned to condensate (spot) data, enabling annotation at both single-cell and well levels. Per-cell features, including condensate count, mean condensate intensity and condensate size distribution, were computed from stringently filtered metadata restricted to double-transfected cells (e.g., PhyB-PIF3). Final outputs were aggregated at the cell, well and condition levels and exported as Excel files for downstream statistical analysis and visualization. All experiments were performed in three independent replicates. An overview of the KNIME workflows is provided here (Fig. S4), while the full workflows, including the details of the consolidated nodes, are available from the authors upon reasonable request.

### Statistical analysis

Statistical analyses were performed using GraphPad Prism (version 10). For experiments evaluating temperature responses, data were fitted with linear or quadratic regression models. When a single model failed to adequately describe the entire temperature range, separate regression lines were fitted for the cold-to-warm and warm-to-hot intervals. The significance of the quadratic regression was evaluated against the linear model, and the linear regression was tested against the null (mean-only) model.

For comparisons involving only two temperature conditions, an unpaired Student’s *t*-test was used. For experiments involving three temperature conditions or factorial combinations of temperature and other treatments (e.g., Picloram, R/FR pulses), one-way or two-way ANOVAs were employed, followed by Tukey’s post-hoc tests for multiple comparisons.

Representative images were selected from the dataset used for quantitative analysis and are displayed side-by-side. The quantitative values of these specific images reflect the mean values for their respective conditions. For confocal microscopy, the observed fluorescence is specific to the indicated probe, as all other signals were filtered out.

The meta-analysis of the expression of genes corresponding to the gene ontology term “response to auxin” (GO:0009733) was conducted using the same procedures and list of published experiments described earlier (Bianchimano *et al*., 2023).

## Results

### Impaired growth inhibition by hot temperatures in auxin-related mutants

We cultivated seedlings of *Arabidopsis thaliana* under short days at 22°C, transferred them to temperatures ranging from 10°C to 37°C 1 h after the beginning of the fourth photoperiod and measured hypocotyl elongation over the next 9 h. Wild-type Col-0 showed a monophasic growth curve peaking at 28°C (Fig. 1a-b). Mutants impaired in auxin biosynthesis, the *shade avoidance 3 (sav3)* and in the *yucca3 (yuc3) yuc5 yuc7 yuc8 yuc9* quintuple (*yucq*), showed reduced elongation at warm temperatures^14,21^ and, strikingly, lacked the growth reduction between 28°C and 37°C (Fig. 1b). In these mutants, growth at cold temperatures was substantially lower than under heat, demonstrating that inhibition was feasible but not observed.

**Figure 1.**
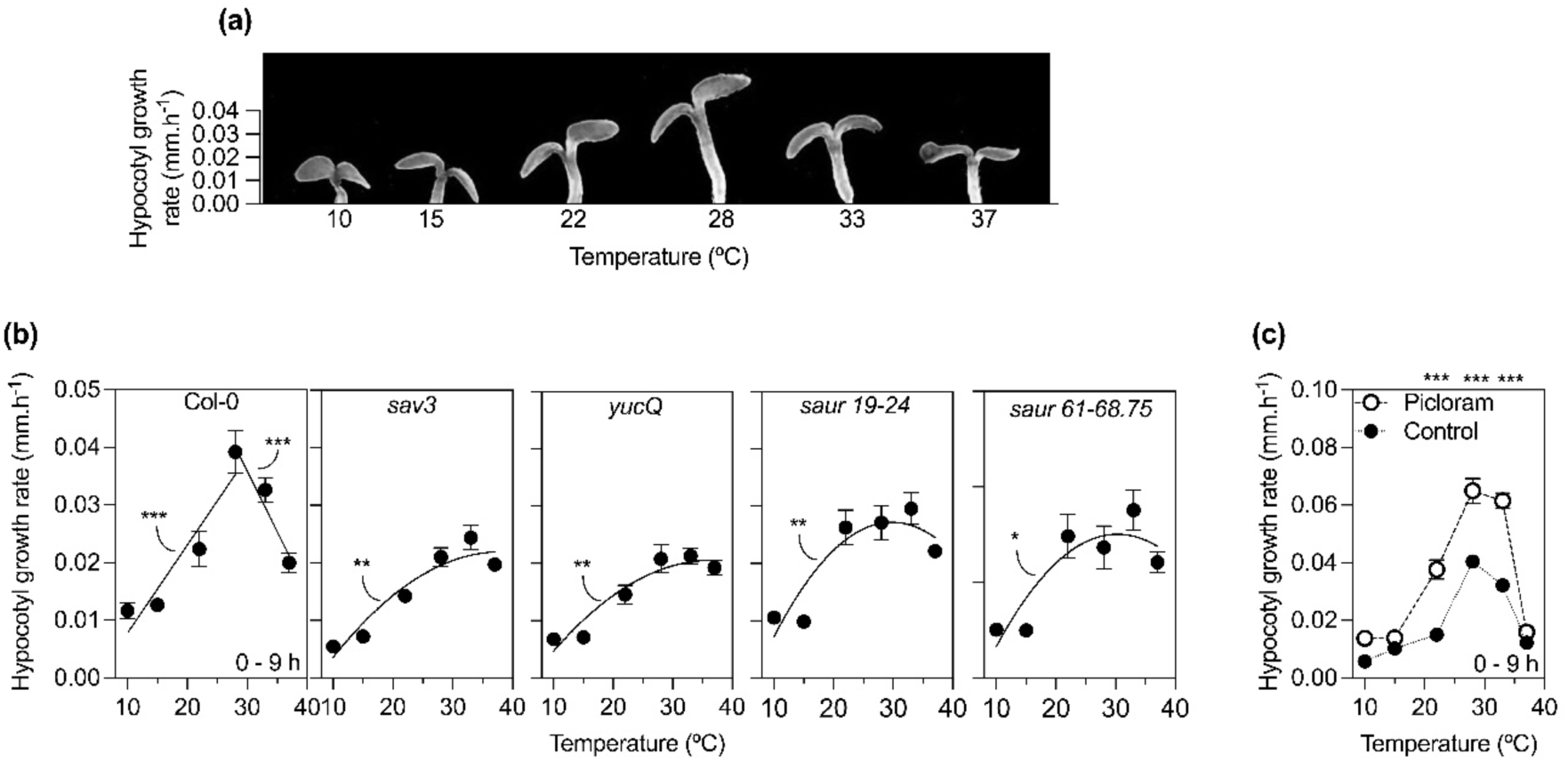
Impaired growth inhibition by hot temperatures in auxin-related mutants. (a) Representative of Col-0 seedlings grown for 9 h at the indicated temperatures. The base of the pictures corresponds to the top of the hypocotyl at the beginning of the treatments. (b) Compared to Col-0 seedlings, the auxin-related *sav3*, *yucQ*, *saur19–24* and *saur61–68, 75* mutants show impaired promotion of hypocotyl growth by warm temperature and lack heat-induced growth inhibition. (c) Exogenous application of the synthetic auxin Picloram (Table S1) promotes hypocotyl growth but does not fully restore heat-inhibited growth. The seedlings were grown at 22°C, transferred to the indicated temperatures 1 h after the beginning of the photoperiod of the fourth day, and photographed 0 and 9 h after the beginning of temperature treatments to measure hypocotyl growth. Each datum point is the mean ±SE of 6 biological replicates pooled from three independent experiments. Asterisks denote statistical significance of linear or quadratic regression lines and of the differences between plants treated with Picloram and controls at a given temperature: ***, P <0.001; **, P <0.01; *, P <0.05.

Auxin promotes the expression of *SMALL AUXIN UP RNA (SAUR)* genes, which help acidify the apoplast to promote cell growth (Spartz *et al*., 2014; Weijers & Wagner, 2016; Vanneste *et al*., 2025). The *saur* multiple mutants (*saur19*-*24* and *saur60-68, 75*) also failed to show growth inhibition by heat compared to warmth (Fig. 1b), supporting the idea that growth inhibition at higher temperatures may result from insufficient, rather than excessive, auxin. Consistently with this interpretation, exogenous auxin partially restored hypocotyl elongation at 33°C (Fig. 1c). However, exogenous auxin did not restore growth at 37°C, suggesting that additional auxin level-independent mechanisms could contribute to heat-induced growth inhibition.

### Hot temperatures reduce auxin abundance and signalling

Prompted by the impaired growth responses to heat observed in the auxin-related mutants, we investigated the status of the canonical nuclear auxin signalling pathway. The levels of the hormone measured in hypocotyl cells with the AuxSen reporter and by a colorimetric method in entire seedlings peaked at 28°C and declined towards 33-37°C (Fig. 2a-b). Auxin signalling involves decreased Aux/IAA co-receptors abundance and increased ARFs transcriptional activity, promoting auxin-responsive gene expression (Weijers & Wagner, 2016; Vanneste *et al*., 2025). Consistently, DII-VENUS reporters indicated minimal Aux/IAA stability at 28°C and higher levels at 33-37°C (Fig. 2c). The *DR5* promoter is a reporter of the transcriptional activity of ARFs and peaked at 28°C, declining at hot temperatures (Fig. 2d-f).

**Figure 2.**
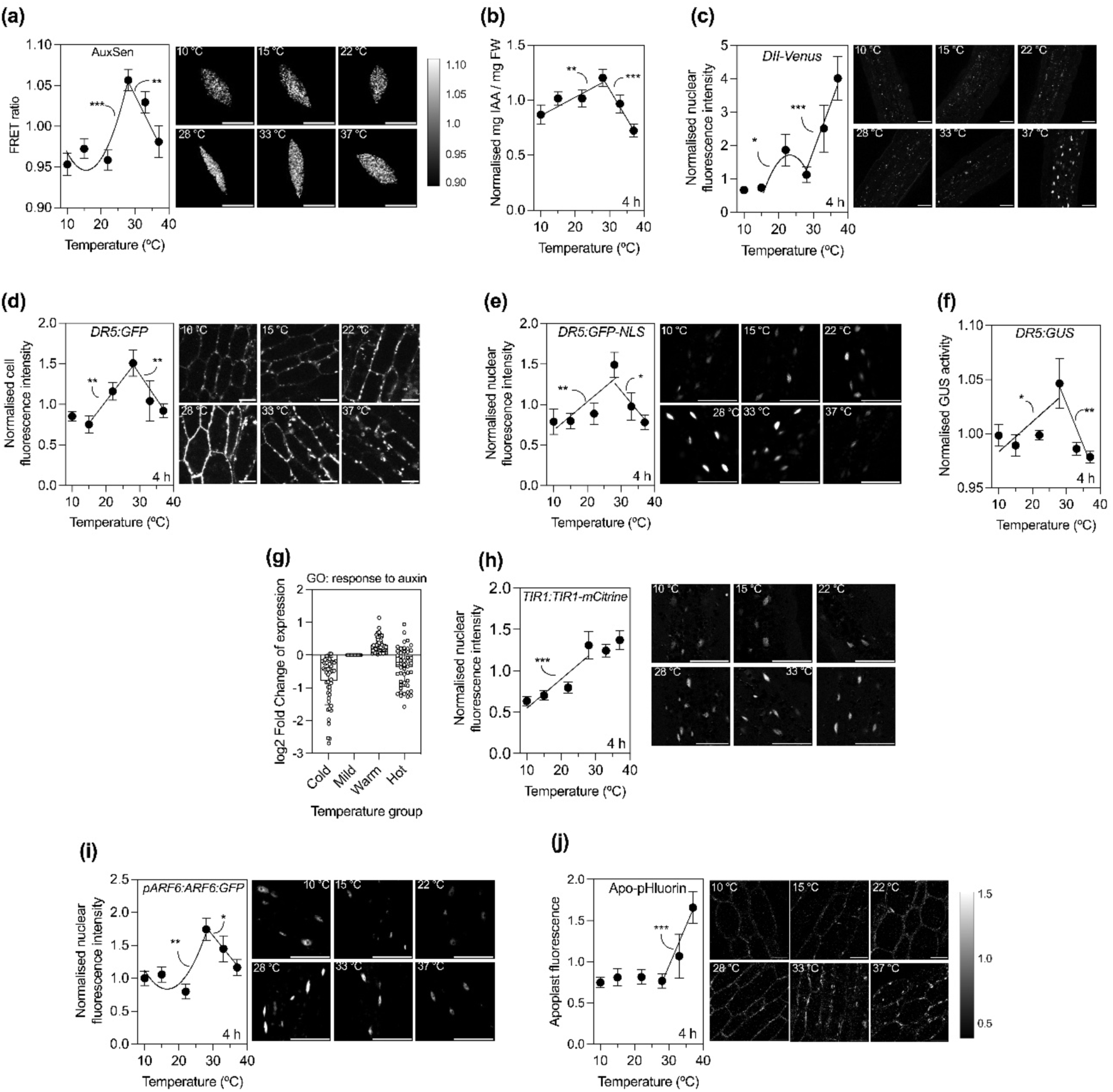
Hot temperatures reduce auxin abundance, perception and signalling. (a-b) Auxin quantification with the FRET sensor AuxSen in hypocotyl cells (a) or by a colorimetric assay in whole seedlings (b) show a peak at warm temperature and a decline at hot temperatures. (c) Confocal analysis of DII-VENUS in hypocotyl cells shows a minimum at warm temperature (indicating high auxin signalling) and an increase at hot temperatures (indicating low auxin signalling). (d-f) Confocal analysis of *DR5:GFP* reporters in hypocotyl cells (d-e, two different reporters) or bioluminescence assays with the *DR5:GUS* reporter (f) indicate maximal ARF transcriptional activity at warm temperatures and reduced activity at hot temperatures. (g) Meta-analysis of the expression of genes corresponding to the GO term “response to auxin” shows a peak at warm temperatures and a decline at hot temperatures. (h) Confocal analysis of TIR1-mCitrine in hypocotyl nuclei indicates increased abundance of the TIR1 auxin receptor at warm temperatures and stable values at hot temperatures. (i) Confocal analysis of ARF6-GFP in hypocotyl nuclei indicates increased abundance of ARF6 transcription factor at warm temperatures and a decrease at hot temperatures. (j) Confocal analysis of the apoplastic pH reporter Apo-pHluorin shows increased pH at hot temperatures. The seedlings were grown at 22°C, transferred to the indicated temperatures 1 h after the beginning of the photoperiod of the fourth day, and harvested 4 h after the beginning of temperature treatments. Scale bars: 5 µm (a) or 50 µm (c-e, g, j, i). Each datum point is the mean ±SE of 12-16 seedlings (a, c-f, h-j) or 3 biological replicates (b), pooled from three independent experiments. Data of meta-analysis (I) are the mean ±SE of log2 Fold Change of expression for 48 genes responsive to auxin (GO:0009733) that responded significantly to temperature. Asterisks denote statistical significance of linear or quadratic regression lines and between control: ***, *P* <0.001; **, *P* <0.01; *, *P* <0.05.

Using a meta-analysis approach previously developed in our group (Bianchimano *et al*., 2023), temperature-responsive genes annotated with “response to auxin” similarly peaked at warm temperatures and declined at hot temperatures (Fig. 2g). The canonical auxin pathway is active at warm temperatures but declines under heat.

Auxin receptor TIR1-mCITRINE fluorescence in hypocotyls increased at 28°C (Wang *et al*., 2016) and remained high at 33-37°C (Fig. 2h). In contrast, nuclear ARF6-GFP levels rose at 28°C (Wilkinson *et al*., 2026) but decreased under heat (Fig. 2i), supporting the above interpretation that signalling components downstream the hormone itself are affected by high temperatures.

Heat-induced growth inhibition was also accompanied by apoplastic alkalization, as indicated by increased fluorescence of the apoplast-targeted pH sensor Apo-pHluorin, which reports extracellular pH with higher fluorescence meaning more alkaline conditions (Fig. 2j). This change is consistent with reduced auxin signalling and opposite to the alternative hypothesis involving excessive auxin, which inhibits growth by causing supra-optimal acidification (Wang *et al*., 2025).

### Functional analysis of key thermomorphogenic genes indicates a reversal of their physiological impact at supra-optimal temperatures

We measured hypocotyl growth in wild-type plants and in mutants of phyB and the PIFs (PIF4, PIF7, PIF5, and PIF3, Fig. 3a). We then used these data to model the contribution of each gene to hypocotyl growth inhibition or promotion under various temperature conditions (see the Methods section for a detailed description of the calculations). As expected, comparing cold and warm temperatures, the inhibitory contribution of phyB decreased, while the promotive contributions of the PIFs increased (Fig. 3b). Surprisingly, this trend reversed at supra-optimal temperatures. The inhibitory activity of phyB was enhanced, whereas the growth-promotion activities of PIF7, PIF4, PIF5 and PIF3 declined (Fig 3b).

**Figure 3.**
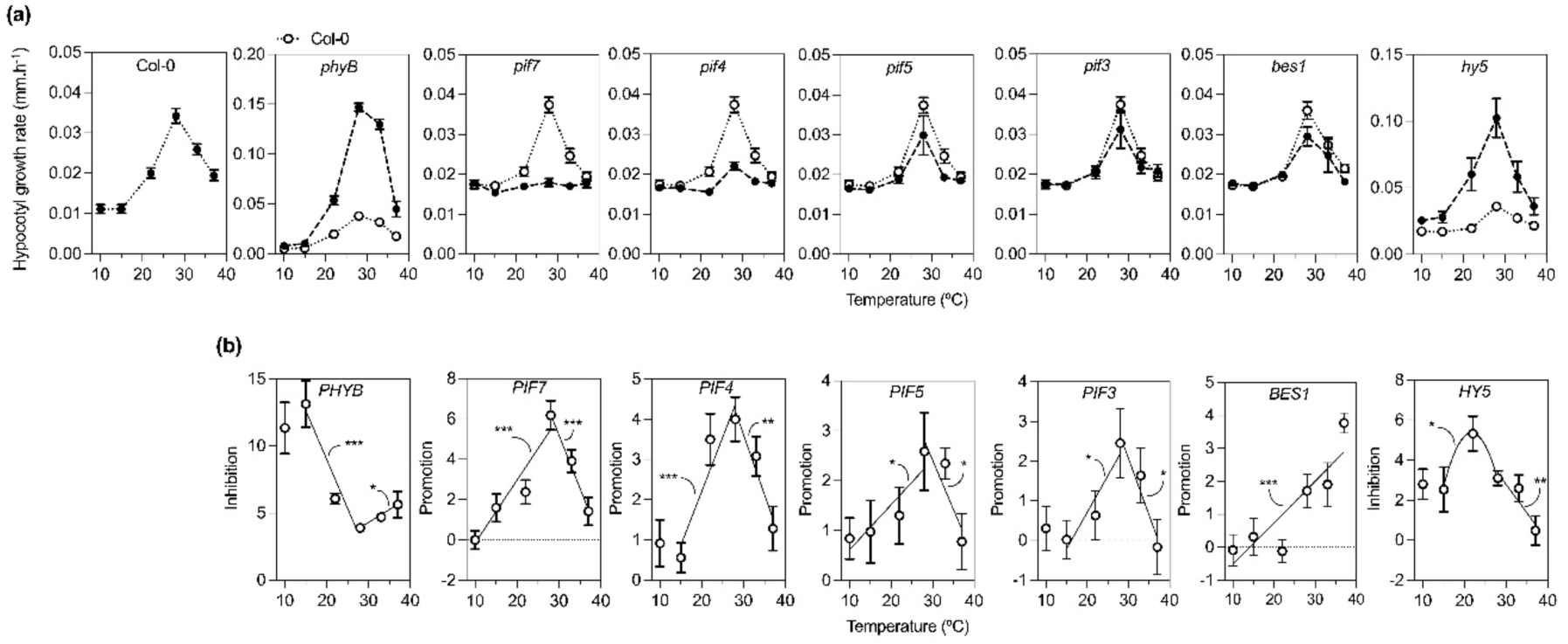
Functional analysis of key thermomorphogenic genes indicates a reversal of their physiological impact at supra-optimal temperatures. (a) Hypocotyl growth of Col-0 wild-type seedlings and *phyB*, *pif7*, *pif4*, *pif5*, *pif3*, *bes1* and *hy5* mutants exposed to different temperatures. (b) Impact of the *PHYB, PIF7, PIF4, PIF5, PIF3, BES1* and *HY5* genes on hypocotyl growth calculated by using a two-equation mathematical model incorporating the wild-type and null mutant data shown in A, as described in the Methods section. The seedlings were grown at 22°C, transferred to the indicated temperatures 1 h after the beginning of the photoperiod of the fourth day, and photographed 0 and 9 h after the beginning of temperature treatments. Each datum point is the mean ±SE of 8-14 biological replicates pooled from three independent experiments. Asterisks denote statistical significance of linear or quadratic regression lines: ***, *P* <0.001; **, *P* <0.01; *, *P* <0.05.

The transcription factor BRI1-EMS-SUPPRESSOR 1 (BES1) promotes hypocotyl growth at warm temperatures by acting directly in the hypocotyl, rather than by influencing auxin synthesis in the cotyledons (Ibañez *et al*., 2018; Costigliolo Rojas *et al*., 2022). Our data confirmed this function, showing an increased promotive contribution between cold and warm temperatures (Fig. 3b).

Notably, BES1 did not show a reversal under heat (Fig. 3b), which aligns with its action outside the main auxin-synthesis organ.

The transcription factor ELONGATED HYPOCOTYL 5 (HY5) inhibits hypocotyl elongation. The observation that *hy5* mutants exhibit hypersensitive hypocotyl growth at warm temperatures (Delker *et al*., 2014) suggests that the inhibitory activity of HY5 is enhanced at warmer temperatures. Our data supports this by showing maximal HY5-mediated inhibition at warm temperatures (Fig. 3b). Importantly, the inhibitory impact of HY5 decreased under heat, mirroring the reduced promotive effect of PIF4 (Fig. 3b), is consistent with the established model that HY5 negatively regulates PIF4 (Johansson *et al*., 2014; Gangappa *et al*., 2017).

### Reduced PIF4 protein abundance in the cotyledons at hot temperatures

Motivated by the results of the genetic analysis, we investigated the abundance of PIF4 which is known to promote auxin synthesis in cotyledons in response to warmth (Franklin *et al*., 2011; Sun *et al*., 2012), The results revealed a complex, organ-specific response.

Using confocal microscopy, we observed that PIF4-GFP protein levels in cotyledons and hypocotyls behaved differently. While the fluorescent signal increased in both organs when the temperature shifted from cold to warm, it showed a distinct pattern at hot temperatures. In hypocotyls, PIF4-GFP continued to rise, whereas in cotyledons, protein levels decreased (Fig. 4a-d). This reduction in cotyledons was more pronounced after 8 h than at 4 h, suggesting a delayed inhibitory effect of high temperature.

**Figure 4.**
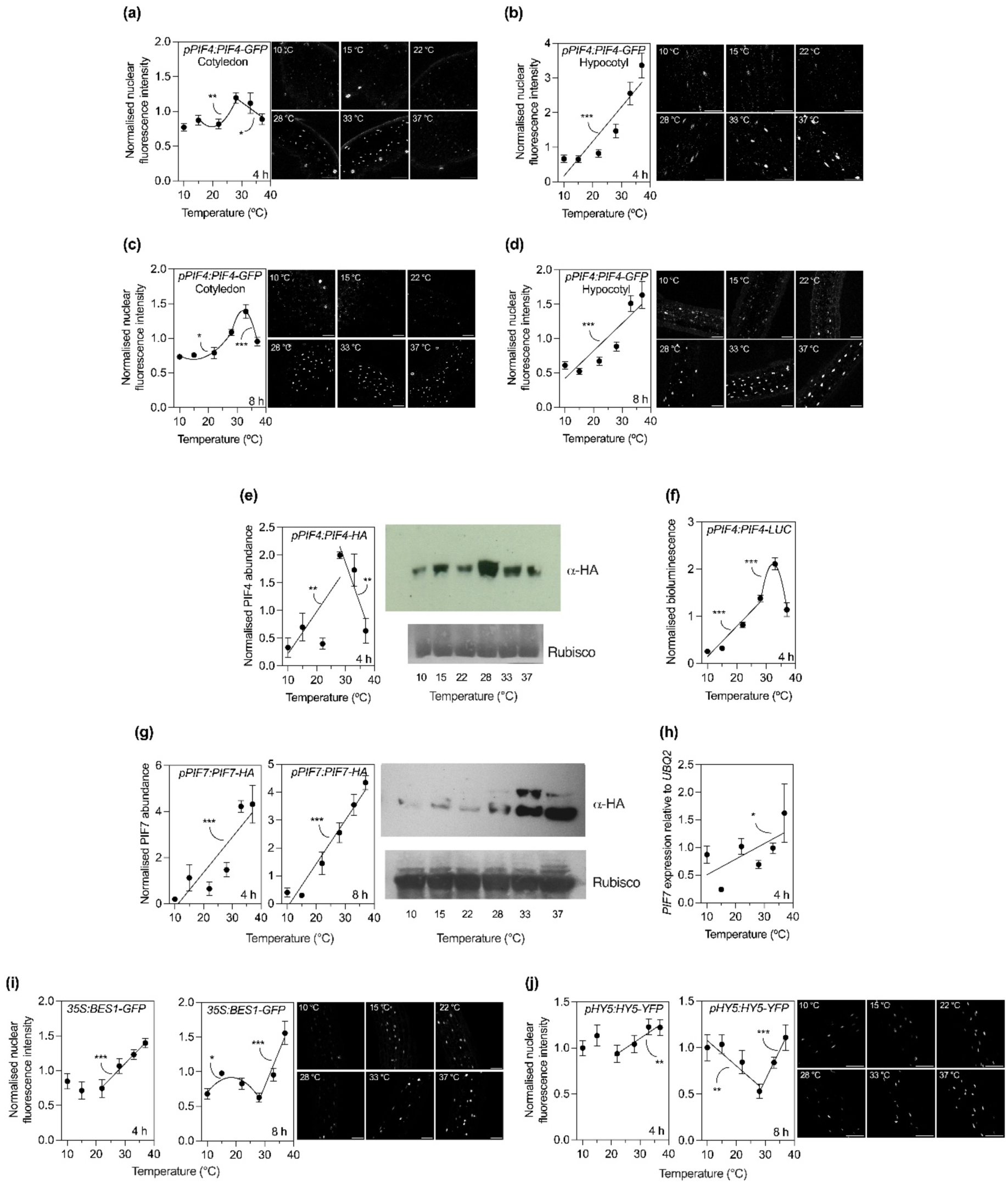
Reduced PIF4 protein abundance at hot temperatures. (a-d) Confocal analysis of the fluorescence driven by the *pPIF4:PIF4-GFP* reporter in the cotyledons (a, c) or the hypocotyl (b, d) in seedlings exposed to different temperatures for 4 h (a, b) or 8 h (c, d). (e) Protein blot of PIF4-HA in seedlings exposed to different temperatures for 4 h. (f) Luminescence driven by the *pPIF4:PIF4-LUC* reporter in seedlings exposed to different temperatures for 4 h. (g) Protein blot of PIF7-HA in seedlings expressing the pPIF7:PIF7-HA transgene and exposed to different temperatures for 4 h or 8 h. (h) Levels of the *PIF7* transcripts measured by qRT-PCR in seedlings exposed to different temperatures for 4 h. (i-j) Confocal analysis of the fluorescence driven by the *p35S:BES1-GFP* reporter (i) or the *pHY5:HY5-YFP* reporter (j) in seedlings exposed to different temperatures for 4 h or 8 h. The seedlings were grown at 22°C, transferred to the indicated temperatures 1 h after the beginning of the photoperiod of the fourth day, and harvested 4 h or 8 h after the beginning of temperature treatments. Scale bars: 50 µm (a-d, i-j). Each datum point is the mean ±SE of 14 seedlings (a-d, i-j) or 3 (e, i), 6 (f), or 4 (g) biological replicates pooled from three (a-d, i-j) or two (e-i) independent experiments. Asterisks denote statistical significance of linear or quadratic regression lines: ***, *P* <0.001; **, *P* <0.01; *, *P* <0.05.

Additional experiments using protein blots and luminometer readings supported these findings. Protein blots of whole seedlings, dominated by cotyledon tissue, showed maximal PIF4 abundance at warm temperatures and, decreased at both temperature extremes, aligning with the cotyledon-specific response (Fig. 4e, Fig. S1). Similarly, luminometer readings from PIF4-LUC, which are also primarily influenced by the cotyledon signal, confirmed a decrease in PIF4 at hot temperatures (Fig. 4f).

### The different behaviour of PIF7, BES1, and HY5

PIF7 is also involved in the promotion of auxin synthesis (Fiorucci *et al*., 2020). However, in contrast to PIF4, its protein levels peaked at the highest temperatures tested in this study (Fig. 4g, Fig. S2), likely driven by enhanced gene expression (Fig. 4h). Similarly, the nuclear abundance of BES1-GFP continued to increase at supra-optimal temperatures (Fig. 4i). Conversely, HY5-YFP levels decreased at warm temperatures, consistent with its known destabilisation (Park *et al*., 2017), but surprisingly recovered under heat stress, exhibiting a response with a minimum signal at intermediate temperatures (Fig. 4j).

While BES1 levels correlated with its physiological activity, the patterns of PIF7 and HY5 did not align with their respective physiological impacts (Fig. 3b). This discrepancy suggests that the activities of PIF7 and HY5 are likely modulated by complex protein-protein interactions or systemic components, rather than being driven solely by their own temperature-dependent protein abundances.

### Increased phyB NB size at hot temperatures

Previous studies indicate that phyB NB size peaks at mild temperatures and decrease at both cold and warm extremes (Legris *et al*., 2016; Murcia *et al*., 2020). Using various phyB-GFP transgenic lines, we confirmed this general trend in hypocotyl cells within the cold-to-warm range; however, we also observed an unexpected increase in NB size at hot temperatures (Fig. 5a-c). This increase at hot temperatures mirrors the physiological activity of phyB estimated from our genetic experiments (Fig. 3b). Notably, these larger heat-responsive NBs were also present in cotyledon cells (Fig. S5).

**Figure 5.**
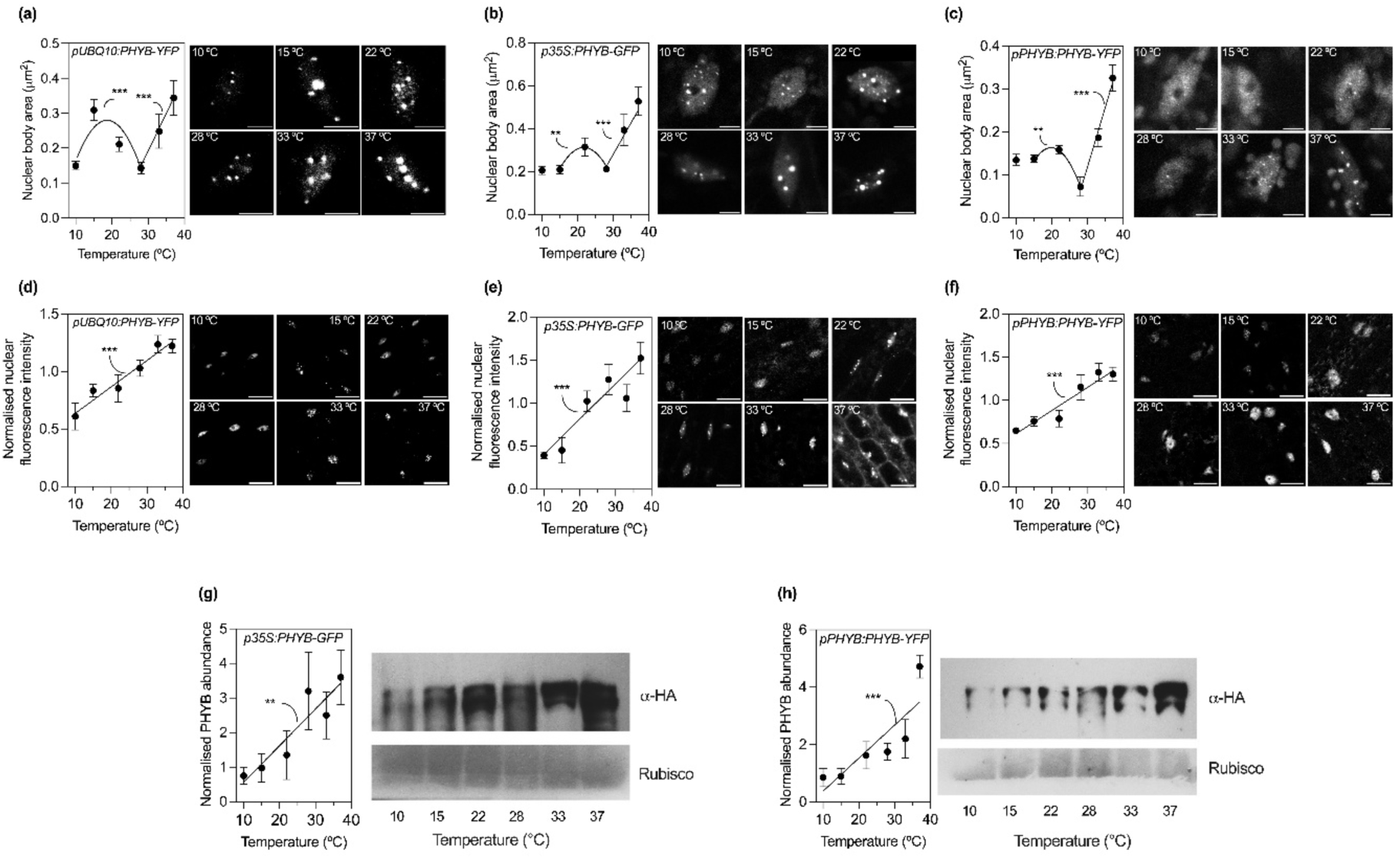
Increased phyB NB size at hot temperatures. (a-c) Confocal analysis of the area of the NBs in hypocotyl cells driven by *pUBQ10:PHYB-YFP* (a), *p35S:PHYB-GFP* (B) or *pPHYB:PHYB-YFP* (c). (d-f) Confocal analysis of the nuclear fluorescence in hypocotyl cells driven by *pUBQ10:PHYB-YFP* (D), *p35S:PHYB-GFP* (E) or *pPHYB:PHYB-YFP* (f). (g-h) Protein blot of PHYB-GFP in seedlings expressing the *p35S:PHYB-GFP* (g) or *pPHYB:PHYB-YFP* (h) transgenes. The seedlings were grown at 22°C, transferred to the indicated temperatures 1 h after the beginning of the photoperiod of the fourth day, and harvested 4 h after the beginning of temperature treatments. Scale bars: 5 µm (a-c) or 20 µm (d-f). Each datum point is the mean ±SE of 12-16 seedlings (a-f) or 4 biological replicates (g-h) pooled from three (a-f) or two (g-h) independent experiments. Asterisks denote statistical significance of linear or quadratic regression lines: ***, *P* <0.001; **, *P* <0.01; *, *P* <0.05.

In addition, phyB-GFP nuclear fluorescence increased across all tested temperatures (Fig. 5d-f), a trend confirmed by protein blots, which showed a corresponding increase in total phyB protein levels (Fig. 5g-h, Fig. S3 see also reference (Hahm *et al*., 2020)). While the decrease in phyB NB size at higher temperatures is known to be caused by thermal reversion of phyB (Legris *et al*., 2016), the increase in their size under hot conditions could be related at least partially to higher levels of nuclear phyB (Ruff *et al*., 2018; Alberti *et al*., 2019; Villegas *et al*., 2022).

### High temperatures promote phyB NB formation despite FR-induced inactivation

Since heat promotes the formation of large NBs while FR triggers their disassembly (Rausenberger *et al*., 2010; Van Buskirk *et al*., 2014), we investigated the outcome of combining these antagonistic stimuli. Arabidopsis seedlings were grown at 22°C and, 1 h after the beginning of the fourth photoperiod, exposed to a 10-minute pulse of either R or FR before transfer to darkness at either 28°C or 37°C.

Seedlings transferred to 28°C for 4 h following an FR pulse exhibited only very small NBs (Fig. 6a). A similar result was observed in seedlings receiving an R pulse. Although R initially establishes a high proportion of the biologically active Pfr form, rapid thermal reversion at 28°C (Legris *et al*., 2016; Burgie *et al*., 2021) gradually depletes this pool, leading to a progressive reduction in NB size(Van Buskirk *et al*., 2014). Supporting this interpretation, NBs persisted at 28°C when R pulses were applied hourly to maintain the active Pfr pool.

**Figure 6.**
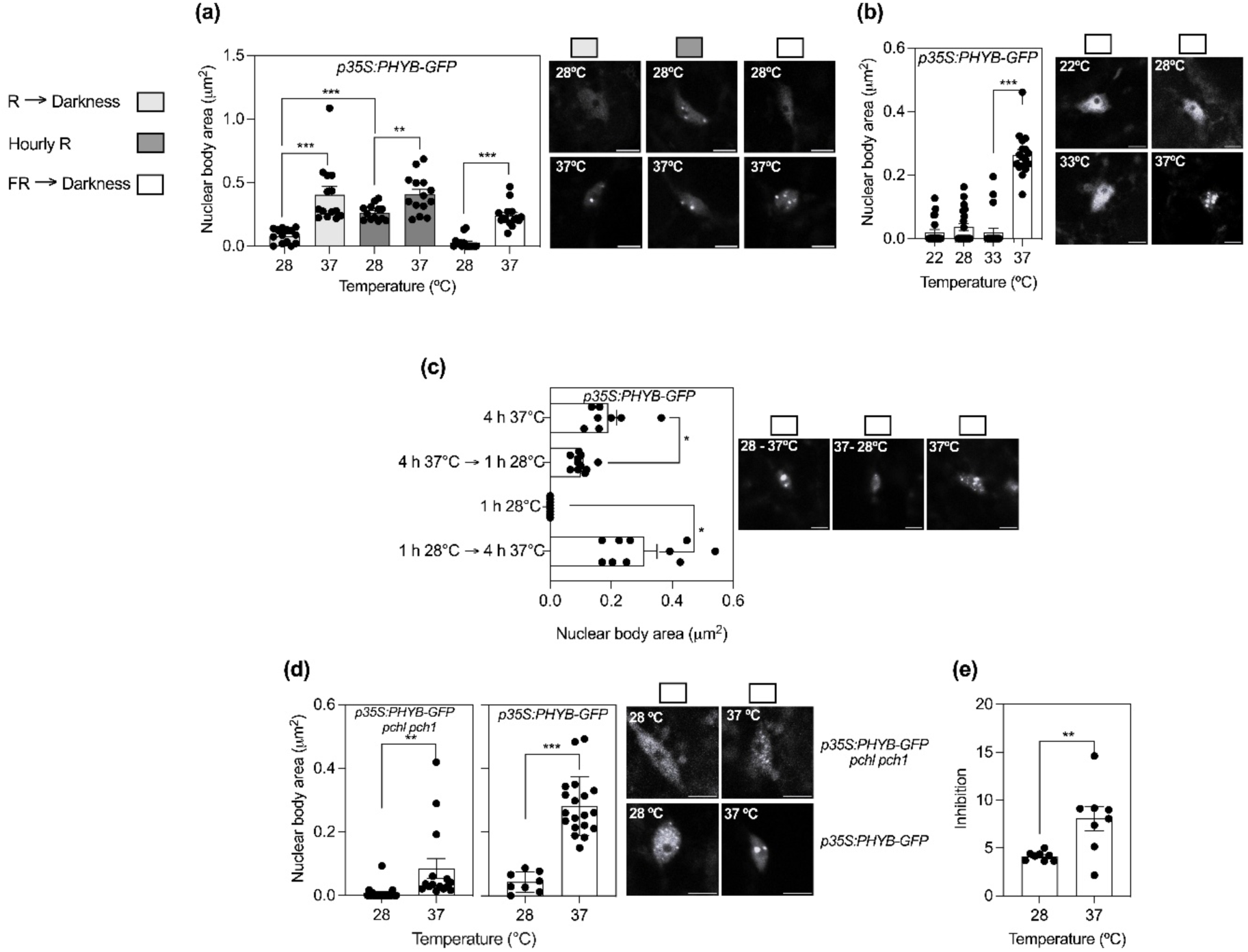
High temperatures promote phyB NB formation despite FR-induced inactivation. (a-d) Confocal analysis of the area of the NBs driven by *p35S:PHYB-GFP* in seedlings exposed to the indicated light and temperature conditions. In D (left) the transgene is expressed in the *pch1 pchl* background. (e) Impact of the *PHYB* gene on hypocotyl growth calculated by using a two-equation mathematical model incorporating the wild-type and *phyB-9* null mutant data, as described in the Methods section. The seedlings were grown at 22°C and, 1 h after the beginning of the fourth day, were transferred to a pulse of R (5 min, 35 μmol m^-2^ s^-1^) followed by darkness, cycles of R (5 min, 35 μmol m^-2^ s^-1^) and darkness (55 min) repeated every hour (the first one immediately after the end of 1 h white light), or a pulse of FR (5 min, 15 μmol m^-2^ s^-1^) followed by darkness. These light / dark conditions were combined with the indicated temperatures. Scale bars: 5 µm (a-d). Data are the mean ±SE and individual values of 8-16 seedlings (a-d) or 8 biological replicates (e), in each case poled from three independent experiments. Asterisks denote statistical significance of ANOVA followed by Tukey tests (a-c) or Student’s t tests (d-e): ***, *P* <0.0001; **, *P* <0.01; *, *P* <0.05.

By contrast, when seedlings were transferred to 37°C, large NBs continued to be present even after an FR pulse (Fig. 6a). This finding is remarkable because the FR pulse converts phyB to its inactive Pr form, which normally promotes NB dissociation (Rausenberger *et al*., 2010; Van Buskirk *et al*., 2014). These results suggest that extreme heat (37°C) exerts a potent effect on phyB NB stability that can bypass the requirement for high levels of active Pfr, although the prior light treatment still influenced the extent of assembly. This effect was specific to extreme heat, as it was observed at 37°C but not at 33°C (Fig. 6b).

To determine whether this thermal effect is reversible, seedlings incubated at 37°C for 4 h following an FR pulse were transferred to 28°C for 1 h. Under these conditions, the NBs formed at 37°C significantly decreased in size (Fig. 6c), indicating that the maintenance of these structures requires sustained high temperature.

Next, we asked whether heat could trigger de novo NB formation. Seedlings treated with a FR pulse were first kept at 28°C for 1 h to allow for near-complete NB disassembly (Fig. 6c). Remarkably, when these seedlings were subsequently transferred to 37°C for 4 h, large phyB NBs successfully reformed (Fig. 6c). These results show that 37°C promote NB assembly even after FR by enhancing phyB nuclear levels and/or phyB propensity to condensate.

We further examined the genetic requirements for this process. In *pch1 pchl* mutant seedlings (Enderle *et al*., 2017; Huang *et al*., 2019) , which strongly impair NB formation, FR treatment followed by darkness yielded NBs approximately ten-fold smaller than in the wild type (Fig. 6d). However, transfer to 37°C significantly increased NB size even in this background (Fig. 6d), further emphasising the potency of the heat-induced assembly mechanism.

Finally, we assessed the physiological relevance of these observations by quantifying hypocotyl growth in wild-type and *phyB* mutant seedlings following a FR pulse and subsequent dark incubation. Using the same approach of Figure 3B, we calculated phyB-dependent growth inhibition. These measurements revealed a clear correlation between heat-induced NB enlargement and enhanced phyB activity (Fig. 6e), suggesting that these assemblies are associated with a functional signalling output. Although phyB activity is typically expected to be low following FR treatment, phyB signalling can be amplified under conditions where only a small Pfr pool remains. For instance, blue light is known to significantly enhance the plant responsiveness to the residual Pfr fraction under FR conditions (Casal & Mazzella, 1998).

### High temperatures promote phyB NB enlargement in an orthogonal mammalian system

To determine whether heat-induced phyB NB formation is dependent on the plant-cell molecular environment, we expressed phyB and its interactor PIF3 in CHO-K1 cells supplemented with a phycocyanobilin chromophore. Following a 15-minute R pulse, cells were incubated in the dark at 22°C, 28°C, or 37°C for 45 min prior to fixation. Consistent with thermal reversion effects, NB size decreased between 22°C and 28°C. Strikingly however, at 37°C, cells exhibited significantly larger NBs (Fig. 7a), resembling the heat-induced NB enlargement observed in Arabidopsis and revealing a biphasic temperature response in which condensates shrink between mild and warm temperatures and re-expand in response to heat. Notably, unlike in plants, this response in CHO-K1 cells did not correlate with significant changes in total nuclear phyB-GFP fluorescence (Fig. 7b), suggesting that heat could increase phyB condensation propensity rather than elevate phyB protein levels.

**Figure 7.**
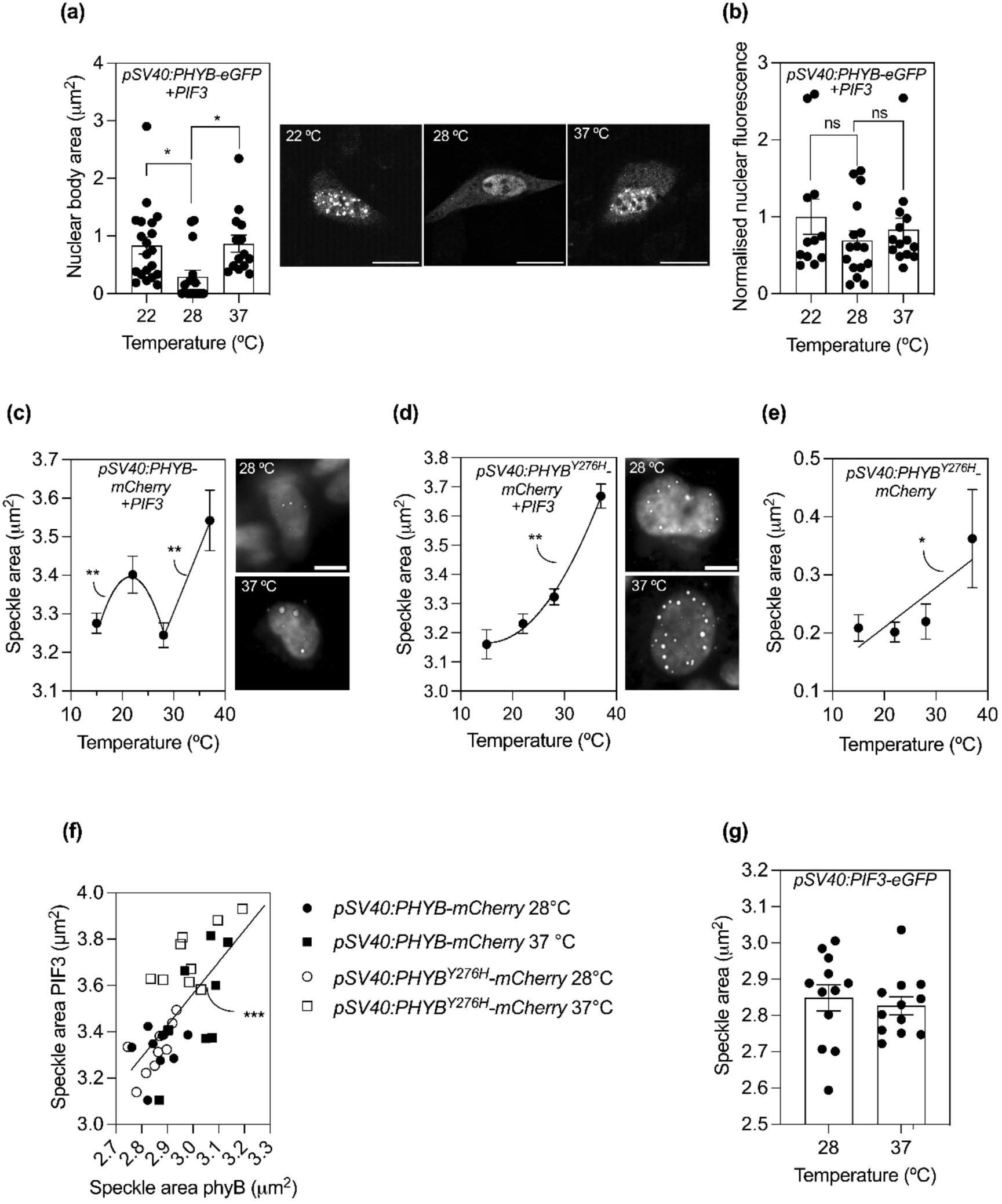
High temperatures enhance phyB condensate size in mammalian cells. (a-b) Confocal analysis of the area of the NBs (a) and total nuclear fluorescence (b) driven by *pSV40:PHYB-eGFP* in CHO-K1 cells co-transfected with *pSV40:PIF3-eCerulean*. (c) Fully automated, high-content quantitative confocal analysis of the area of the speckles driven by *pSV40:PHYB-mCherry* co-transfected with *pSV40:PIF3-eGFP.* Both, nuclear and cytoplasmatic speckles are included in the analysis. (d-e) Fully automated, high-content quantitative confocal analysis of the area of the speckles of *pSV40:PHYB^Y276H^-mCherry* in CHO-K1 cells co-transfected with *pSV40:PIF3-eGFP* (d) or single transfected with *pSV40:PHYB^Y276H^-mCherry* (e). (f) Correlation between the size of the speckles driven by *pSV40:PIF3-eGFP* and the speckles driven by either *pSV40:PHYB-mCherry* or *pSV40:PHYB^Y276H^-mCherry* co-transfected in CHO-K1 cells. (g) The size of the speckles driven by *pSV40:PIF3-eGFP* do not increase in response to heat in single transfected cells. CHO-K1 cells were exposed to 15-min R followed by 60 min of darkness at the indicated temperatures. Scale bars: 20 µm (a). Each datum point is the mean ±SE of 8–10 biological replicates (c-e), the mean ±SE and individual values of 12-20 biological replicates (a-b, g), or individual values (f) pooled from two (a-c, g) or three (d-e) independent experiments. Please refer to Figure S4 for an example showing the number of cells analysed in an individual experiment contributing to c-e. Asterisks denote statistical significance of ANOVA followed by Tukey tests (a-b) or of linear or quadratic regression lines (c-g): ***, *P* <0.0001; **, *P* <0.01; *, *P* <0.05.

The speckles formed by phyB when co-transfected with either PIF4 or PIF7 were smaller and less amenable to quantitative analysis than those observed with PIF3, but an increase in phyB speckle size in response to heat was still detectable for phyB combinations with either PIF4 or PIF7 (Fig. S6).

### High-content automated quantitative analysis of phyB speckle response to temperature

We validated our findings using a fully automated, high-content quantitative approach (Fig. S4). This methodology has two significant advantages for the study of complex patterns because automation minimises the possibility of operator biases and the high-content approach allows for the simultaneous analysis of thousands of CHO-K1 cells. The results confirmed an increase in condensate size at 37°C (Fig. 7c).

To investigate whether this temperature-dependent assembly requires thermal reversion, we analysed the constitutively active phyB^Y276H^ mutant, which lacks thermal reversion (Su & Lagarias, 2007; Hu *et al*., 2009; Wang *et al*., 2024). When co-expressed with PIF3, the size of phyB^Y276H^ condensates also increased with temperature indicating that the enlargement observed at 37°C cannot be explained simply by altered thermal reversion dynamics (Fig. 7d). In CHO-K1 cells single-transfected with phyB^Y276H^ (no PIF3 co-expression), the speckles also increase in size with temperature (Fig. 7e), suggesting this behaviour is likely an intrinsic property of phyB.

We next examined PIF3 speckles that co-localise with either phyB or phyB^Y276H^. These speckles increased in size between 28°C and 37°C, showing the same behaviour as phyB speckles (Fig. 7f). Conversely, PIF3 speckles did not enlarge under heat when phyB was absent (single transfected cells) (Fig. 7g), indicating that PIF3 is not the driver of the response to heat.

### Genetically impaired phyB NB formation reduces growth inhibition under heat

The results above demonstrate that NB enlargement and hypocotyl growth inhibition co-occur under heat stress. To move beyond correlation, we investigated whether genetically reducing NB size lowers growth inhibition by comparing hypocotyl elongation in the wild type against single *pch1*, *pchl* and double *pch1 pchl* mutants exposed to heat. The *pch1 pchl* double mutation reduced NB formation in seedlings exposed to high-irradiance R at 37°C (Fig. 8a). In seedlings exposed to a FR pulse to minimise phyB activity followed by darkness at 28°C, hypocotyl growth was unaffected by the mutations (Fig. 8b). However, under high-irradiance R (maximising phyB activity) at 37°C, *pch1*, *pchl*, and *pch1 pchl* seedlings elongated significantly more than the wild type (Fig. 8b).

**Figure 8.**
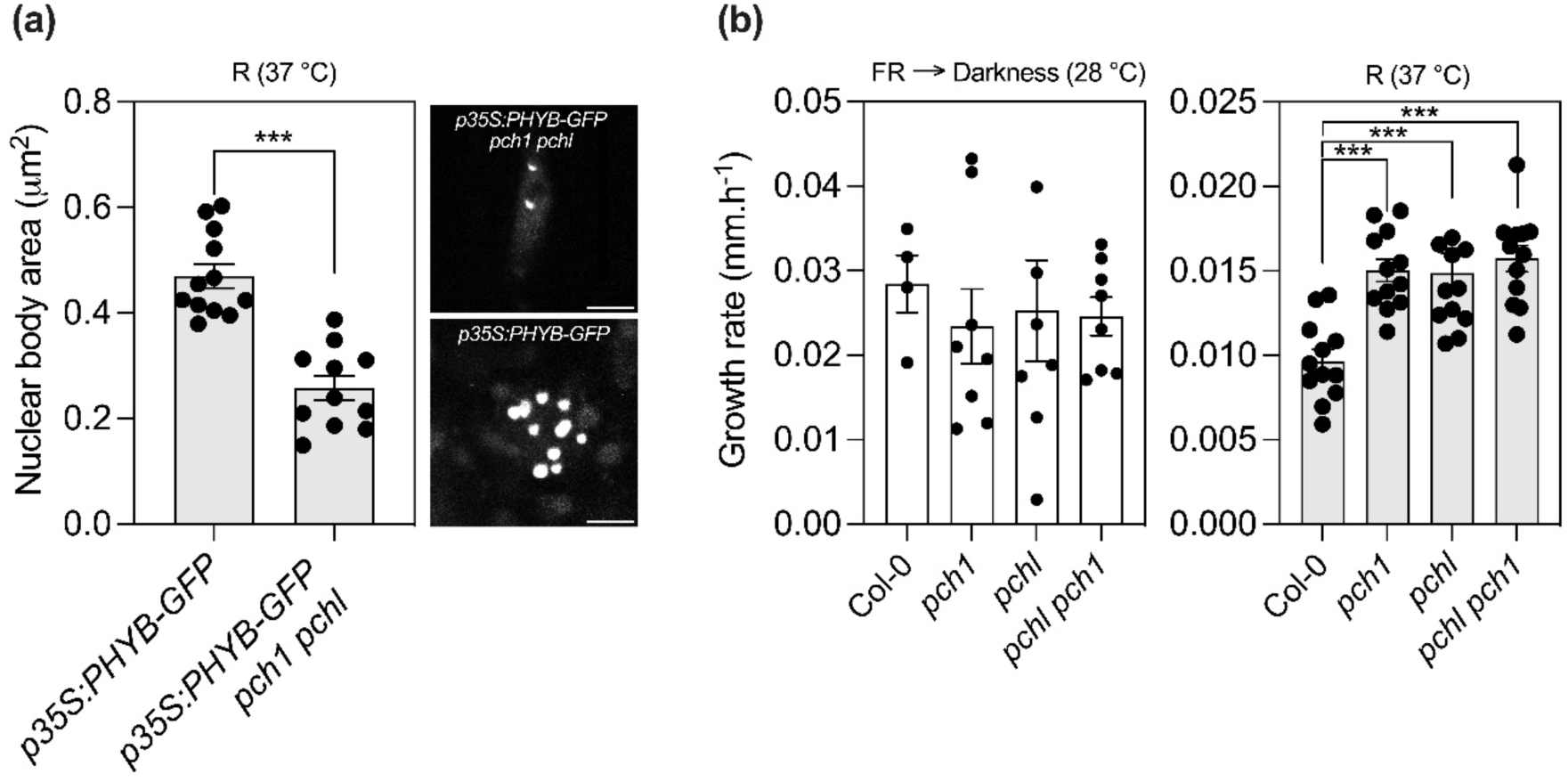
Genetically impaired phyB NB formation reduces growth inhibition under heat. (a) Confocal analysis of the area of the NBs driven by *p35S:PHYB-GFP* in seedlings of the wild-type and *pch1 pchl* mutant backgrounds exposed to high-irradiance continuous R (175 μmol m^-2^ s^-1^) at 37°C. (b) Hypocotyl growth in seedlings of the wild-type and *pch1 pchl* mutant exposed to FR followed by darkness at 28°C (as a control showing no effects of the mutations under conditions of minimal phyB activity) or high-irradiance continuous R at 37°C. Note different scales. The seedlings were grown at 22°C and, 1 h after the beginning of the fourth day, were transferred to a pulse of FR (5 min, 15 μmol m^-2^ s^-1^) followed by darkness at 28°C or high-irradiance continuous R (175 μmol m^-2^ s^-1^) at at 37°C. Scale bars: 5 µm (a). Data are the mean ±SE and individual values of 4-12 biological replicates (a) or 11-12 seedlings (b), in each case pooled from two independent experiments. Asterisks denote statistical significance of ANOVA followed by Tukey tests (a) or Student’s t tests (b): ***, *P* <0.0001.

Establishing direct causality in condensate studies is inherently challenging, as it is "often difficult (if not impossible) to find perturbations that only affect the condensate-forming behaviour while sparing all other functions of the macromolecules involved" (Alberti *et al*., 2025). PCH1 and PCHL are known to exert three primary effects: stabilising the phyB Pfr form by reducing thermal reversion, enhancing phyB NB formation (even in the absence of thermal reversion), and facilitating the degradation of PIF1 protein (Enderle *et al*., 2017; Huang *et al*., 2019; Cheng *et al*., 2020). Notably, these mutations did not exhibit constitutive hypocotyl growth phenotypes in the absence of phyB activation (e.g., through PIF1-mediated effects), as confirmed by our controls exposed to FR at 28°C (Fig. 8b). Furthermore, we minimised the influence of thermal reversion by using high-irradiance continuous R (Burgie *et al*., 2021; Yi *et al*., 2024; Fan *et al*., 2025) ensuring that rapid photochemical reactions outweighed the impact of reversion. Consequently, the increased hypocotyl elongation observed in *pch1* and *pchl* mutants under these specific conditions can be directly attributed to impaired NB formation (Fig. 8a-b).

## Discussion

While plant growth scales with temperature within permissive ranges, our study demonstrates that rapid hypocotyl growth inhibition under heat stress is not a passive consequence of cellular damage, such as protein denaturation or membrane deterioration. Rather, this inhibition results from a coordinated reversal of the phyB–PIF4–auxin pathway that drives growth during cold-to-warm transitions (Quint *et al*., 2023). This regulatory shift is marked by increased phyB abundance and the formation of distinct heat-induced phyB condensates, a reduction in cotyledonary PIF4 levels, decreased auxin levels, and elevated apoplastic pH, which together reinforce growth inhibition (Figs. 1–5).

The regulatory impact of heat is remarkably selective. For instance, while PIF4 levels decreased in the cotyledons (the primary site of auxin synthesis) they continued to rise with temperature in the hypocotyl itself (Fig. 4a-d). Despite this local increase, PIF4 activity in the hypocotyl may still be constrained by phyB-mediated sequestration (Park *et al*., 2018; Pham *et al*., 2018) and a loss of synergism with ARF6 (Oh *et al*., 2014), the levels of which declined under heat (Fig. 2I). In contrast, BES1 (a known component of heat tolerance (Albertos *et al*., 2022)) continued to accumulate and promote growth (Fig. 4I, 3B), highlighting that the heat-induced inhibitory response does not target all growth-promoting pathways equally. Furthermore, levels of the growth-inhibitor HY5 increased under heat, though its maximum physiological effectiveness was observed at lower temperatures (Fig. 4j, 3b). Given that HY5 negatively modulates PIF4 activity(Johansson *et al*., 2014; Gangappa *et al*., 2017), its overall physiological impact likely depends on the stoichiometric interplay between its own abundance and that of its target, PIF4. Similarly, although heat did lower PIF7 protein abundance (Fig. 4g), the reduction in its physiological activity (Fig. 3) likely stems from the depletion of PIF4 precluding the formation of functional PIF4-PIF7 heterodimers(Fiorucci *et al*., 2020) and/or its phosphorylation and sequestration by phyB (Huang *et al*., 2018; Ling *et al*., 2023).

The increased abundance and condensation of phyB (Fig. 5) provide a feasible explanation for its enhanced physiological activity under heat (Fig. 3). Condensation may facilitate the activity of a protein by increasing the "dwelling time" with its partners (Ladbury *et al*., 2023). In support of a role of heat-induced condensates, mutations in phyB interactors that impair NB formation result in longer hypocotyls even when combined with a phyB mutation that adopts a constitutively active conformation (Galvão *et al*.; Enderle *et al*., 2017; Huang *et al*., 2019; Fang *et al*., 2022). More importantly, we observed reduced NB formation and phyB-mediated inhibition of hypocotyl growth in *pch1 pchl* seedlings under heat (Fig. 8). Overexpression of phyB inhibits hypocotyl growth (Wagner *et al*., 1991), suggesting that the higher phyB protein levels also favour the enhanced phyB-mediated growth inhibition under heat.

The distinct nature of the heat-induced phyB NBs is highlighted by the fact that it that they formed even following a FR pulse or in a *pch1 pchl* mutant background (Fig. 6). Because protein concentrations exceeding a critical threshold can trigger the formation of larger condensates via phase separation (Alberti *et al*., 2019), rising phyB levels under heat likely favours this response. However, protein abundance is not the sole driver; heat also appears to enhance the intrinsic propensity of phyB to condense. This was confirmed in mammalian cell, where heat-induced speckle formation occurred even without an increase in phyB protein levels (Fig. 7a–c). In contrast to the decay caused by warm temperatures, the heat-induced increase in phyB NB size does not depend on the occurrence of thermal reversion (Fig. 7d).

The inhibitory responses to heat observed here cannot be explained by a simple, linear transmission of a single temperature input through the phyB-PIF4-auxin module. For instance, treatment at 37°C triggered a decay in ARF6 levels (Fig. 2i), an effect not typically considered a consequence of enhanced phyB activity (Pucciariello *et al*., 2018). Furthermore, warm temperatures influence PIF4 abundance through pathways independent of phyB (Jung *et al*., 2020), which may also contribute to the observed growth inhibition. These results suggest that the system integrates additional, parallel heat-sensing inputs beyond the canonical phyB-PIF4-auxin regulatory axis.

Ecologically, promoting hypocotyl growth at warm temperatures serves as a "foraging" strategy to optimise light capture by outgrowing competing vegetation (Romero-Montepaone *et al*., 2021). However, under extreme heat, this strategy becomes increasingly hazardous. By repressing stem elongation, the plant avoids emerging from the protective shade of the surrounding canopy. This "umbrella effect", where the seedling remains shielded by the foliage of taller neighbours, helps maintain the plant tissue temperature within a tolerable range and reduce water loss, conferring a vital survival advantage and enhancing resilience under severe thermal stress (Miller *et al*., 2021).

## Acknowledgments

Supported by *Agencia Nacional de Promoción Científica y Tecnológica* (ANPCYT, grant PICT-2021-I-A-01070 to JJC), University of Buenos Aires (grant 20020170100505BA to JJC) and *Fundación Instituto Leloir* (institutional grant to JJC). LB received a fellowship from Deutscher Akademischer Austauschdienst (DAAD) to conduct work with mammalian cells in Düsseldorf, Germany, and the Deutsche Forschungsgemeinschaft (DFG, German Research) under the Germany Excellence Strategy (CEPLAS-EXC-2048 project no. 390686111) to M.D.Z. We thank Stefanie Weidtkamp-Peters (Center for Advanced Imaging (CAi) at Heinrich-Heine-University Düsseldorf) for providing access to the Revvity Operetta CLS high-content screening automated microscope system (DFG-INST 208/760-1 FUGG), Stefanie Kuschel for mammalian cell culture technical support, and Hannes M. Beyer and Jonas Schön (Institute of Synthetic Biology, Heinrich-Heine-University Düsseldorf) for helpful discussions.

During the preparation of this work, the authors used Google Gemini to improve the choice of words. The authors reviewed and edited the output as needed and take full responsibility for the content of the published article.

## Author contributions

Investigation, L.B., C.S.K., M.B.D.-L., G.M. and M.E.P.; conceptualization, J.J.C.; writing, J.J.C. with input from L.B., C.S.K., M.B.D.-L., G.M., M.E.P., A.B.H, and M.D.Z. ; funding acquisition, M.D.Z., and J.J.C., visualization, L.B., C.S.K. and J.J.C.; formal analysis, L.B., C.S.K., A.B.H. and J.J.C.; supervision, M.D.Z., and J.J.C.

