## Supporting information for "Thermosensory pathway makes a U-turn to respond to heat"

**Table S1.** Biological materials and reagents used in this study

| **Strains of *Arabidopsis thaliana*** | **Source** | |
|---|---|---|
| **Wild type and mutants** | | |
| Col-0 | N/A | |
| *sav3-1* | (Tao *et al.*, 2008) | |
| *yuc3 yuc5 yuc7 yuc8 yuc9* | (Tao *et al.*, 2008) | |
| *saur 61-68.75* | (Nagpal *et al.*, 2022) | |
| *saur 19-24 #12 and #17* | (Wang *et al.*, 2020) | |
| *phyB-9* | (Reed *et al.*, 1993) | |
| *pif7-1* | (Leivar *et al.*, 2008) | |
| *pif4-101* | (Lorrain *et al.*, 2008) | |
| *pif5-3* | (Fujimori *et al.*, 2004) | |
| *pif3-3* | (Monte *et al.*, 2004) | |
| *bes1-1* | (He *et al.*, 2005) | |
| *hy5-221* | (Shin *et al.*, 2007) | |
| **Reporter lines** | | |
| *AuxSen* | (Herud-Sikimić *et al.*, 2021) | |
| *p35S:DII-Venus* | (Brunoud *et al.*, 2012) | |
| *pDR5:GFP-NLS* | (Liao *et al.*, 2015) | |
| *pDR5:GFP* | (Liao *et al.*, 2015) | |
| *pDR5:GUS* | (Windels *et al.*, 2014) | |
| *pTIR1:TIR1-mCitrine tir1-10* | (Prigge *et al.*, 2020) | |
| *pARF6:ARF6-GFP* | (Rademacher *et al.*, 2011) | |
| *Apo-pHluorin* | (Martinière *et al.*, 2011) | |
| *pPIF4:PIF4-GFP pif4-101* | (Pucciariello *et al.*, 2018) | |
| *pBES1:BES1-GFP* | (Yin *et al.*, 2002) | |
| *pHY5:HY5-YFP* | (Oravecz *et al.*, 2006) | |
| *pARF6:ARF6:GFP* | (Rademacher *et al.*, 2011) | |
| *pPHYB-PHYB-YFP* (*phyB-9*) | (Viczián *et al.*, 2020) | |
| *pUBQ10:PHYB-YFP* (*phyB-9*) | (Zhang *et al.*, 2013) | |
| *p35S:PHYB-GFP phyA-211 phyB-9* | (Enderle *et al.*, 2017) | |
| *p35S:PHYB-GFP phyA-211 phyB-9 pch1 pchl* | (Enderle *et al.*, 2017) | |
| *pPIF4:PIF4-LUC* | (Murcia *et al.*, 2022) | |
| *pPIF4:LUC* | (Murcia *et al.*, 2022) | |
| *pPIF4:PIF4-HA pif4-101 pif5 (pil6-1)* | (Martínez *et al.*, 2018) | |
| *pPIF7:PIF7-3HA-tPIF7 pif7-2* | (Galvão *et al.*, 2019) | |
| **Oligonucleotides for RT-qPCR** | **Sequence 5' >>> 3'** | |
| *PIF7* forward | CGGAGCTTGAAGACAGCTAGAACC | |
| *PIF7* reverse | GACTCGTTGTGAATCGCTGCTG | |
| *UBQ2* forward | AGAATGCTTGGAGTCCTGCTT | |
| *UBQ2* reverse | ACATTTGTGCCATTGAATTGAACCC | |
| **Blot reactives** | **Source** | **Catalog number** |
| HA Tag Polyclonal Antibody (rabbit) | Thermo scientific | PA1985 |
| Anti-rabbit IgG (H+L) HRP Secondary Antibody (goat) | Thermo scientific | 32460 |
| Anti-GFP (mouse) | Roche | 11063100 |
| Anti-mouse (H+L) (goat) | Life Technologies | G21040 |

| | | |
|---|---|---|
| Nitrocellulose Membrane 0.45 µm | Thermo scientific | 88018 |
| **Hormones** | **Source** | **Catalog number** |
| IAA | Sigma-Aldrich | I2886 |
| Picloram | Sigma-Aldrich | P5575 |
| **Cell lines** | **Source** | **Identifier** |
| Hamster: CHO-K1 cells | DSMZ https://www.dsmz.de/collection/catalogue/details/culture/ACC-110M | ACC 110 |
| **Recombinant DNA** | **Source** | **Identifier** |
| *pSV40:PHYB-eGFP* | This paper | pLuB001 |
| *pSV40:PIF3-mCerulean* | This paper | pTB024 |
| *pSV40:PIF4-mCerulean* | This paper | pLuB004 |
| *pSV40:PIF7-mCerulean* | This paper | pLuB006 |
| *pSV40:PIF3-eEGFP* | (Beyer *et al.*, 2015) | pMZ725 |
| *pSV40:PHYBY$^{276H}$-mCHERRY* | This paper | pCSK208A |
| *pSV40:PHYB-mCherry* | (Beyer *et al.*, 2015) | pSJ025 |

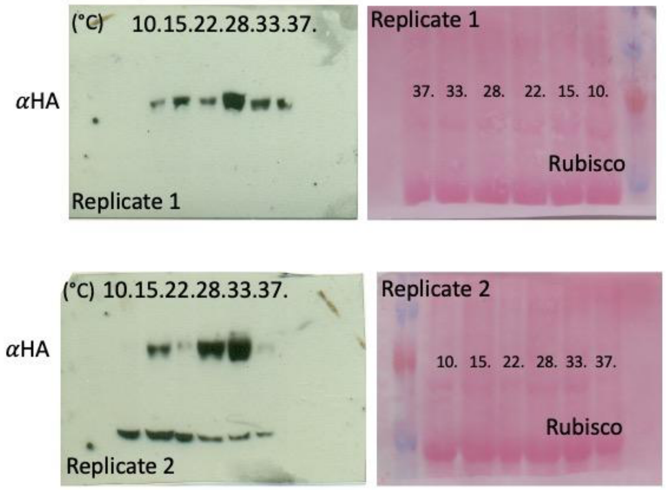


**Fig. S1.** Protein blots of PIF4-HA.

Negative images of the full blots corresponding to each one of the biological replicates (1-2) The same blot was revealed with anti-HA (left) antibodies and with Ponceau staining (right). Temperatures are indicated in °C. Related to Figure 4E and Dataset 1.

4 h

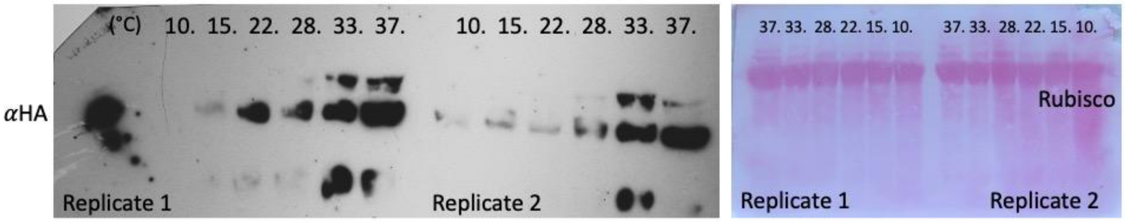


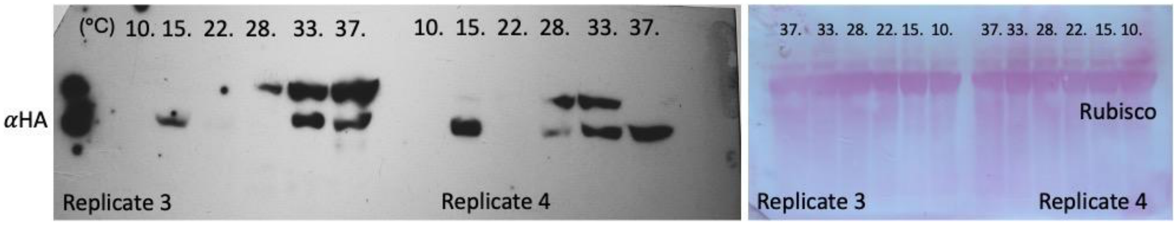


8 h

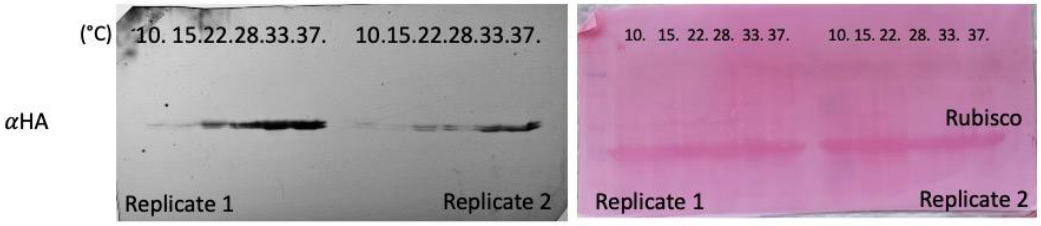


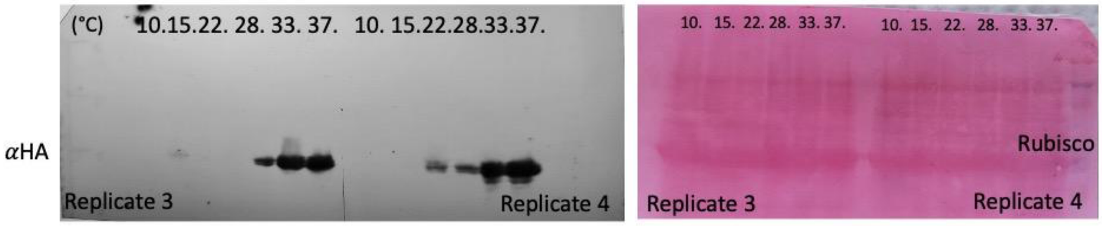


**Figure S2. Protein blots of PIF7-HA.**

Negative images of the full blots corresponding to each one of the biological replicates (1-4 for each time of harvest). The same blot was revealed with anti-HA (left) antibodies and with Ponceau staining (right). Temperatures are indicated in °C. Related to Figure 4G and Dataset 1.

phyB-GFP

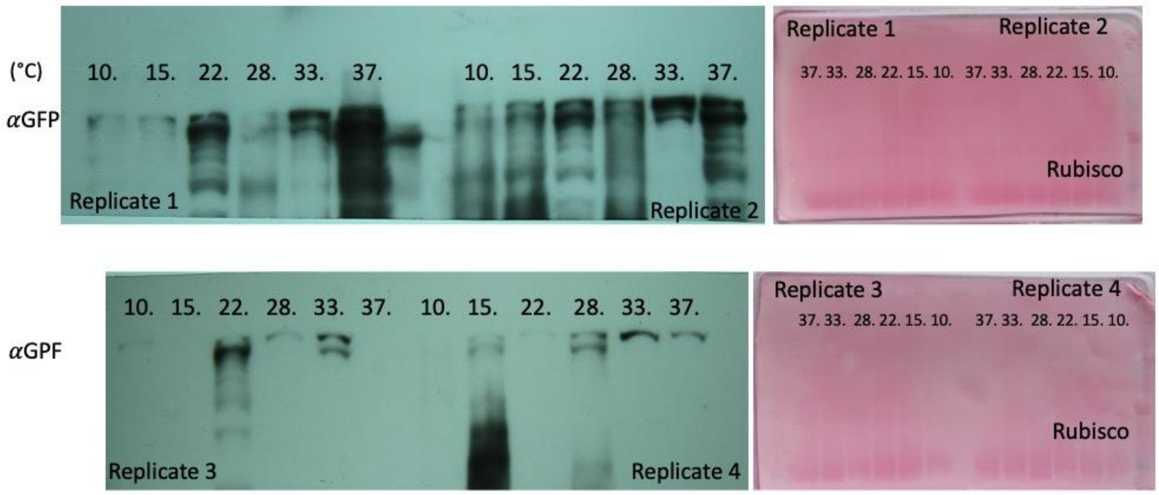


phyB-YFP

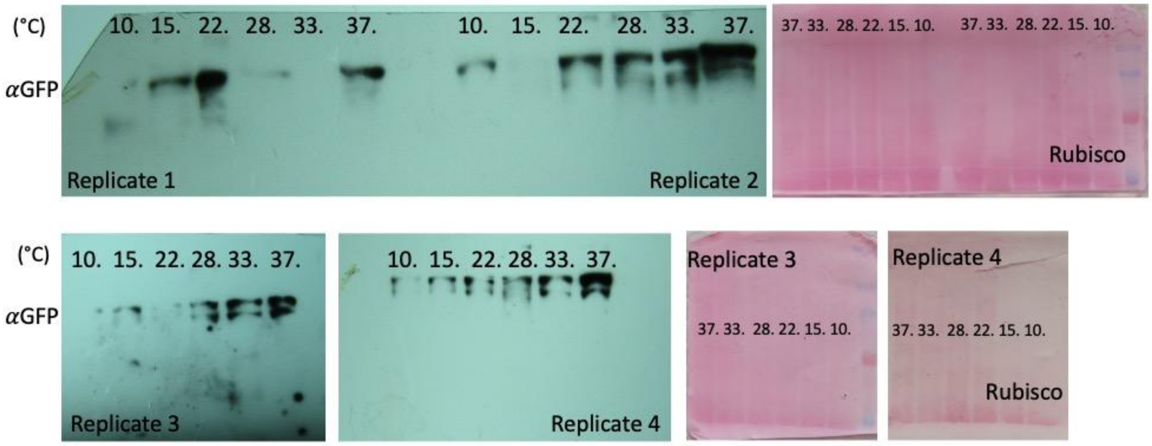


**Figure S3. Protein blots of phyB-GFP and phyB-YFP.**

Negative images of the full blots corresponding to each one of the biological replicates (1-4 for each transgene). The same blot was revealed with anti-GFP (left) antibodies and with Ponceau staining (right). Temperatures are indicated in °C. Related to Figure 5G-H and Dataset 1.

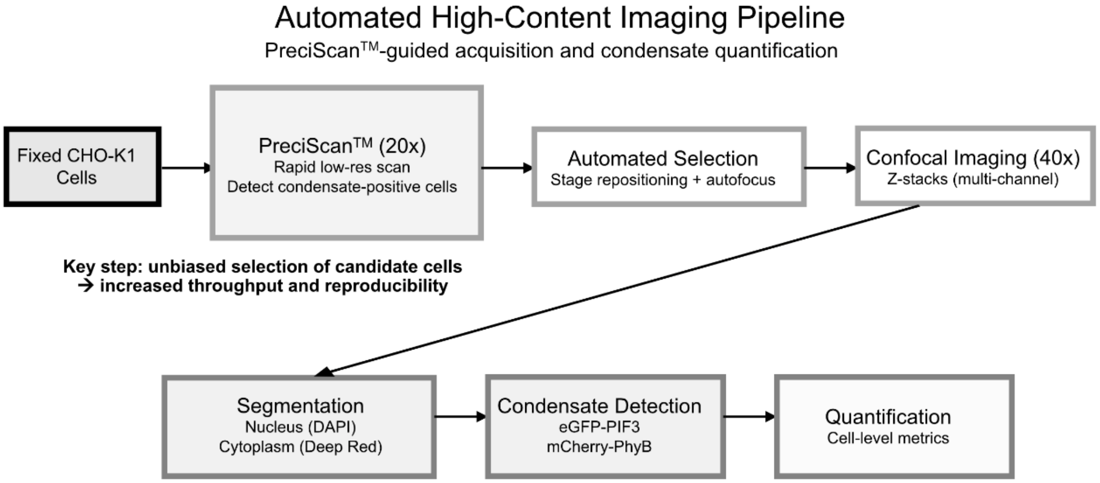


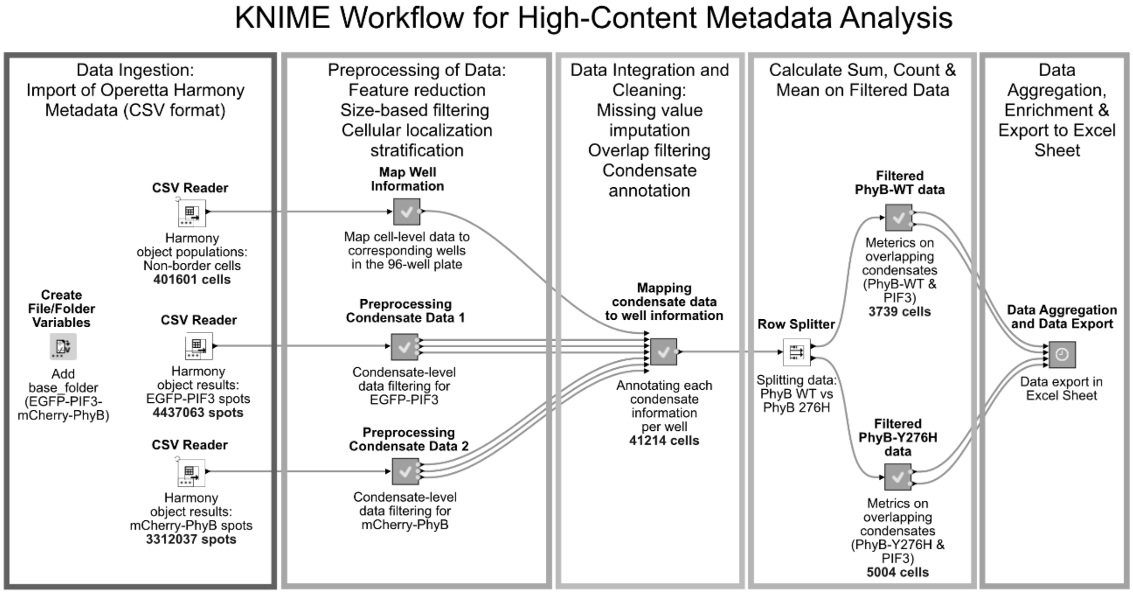


**Figure S4. High-content imaging and analysis of condensates in mammalian cells.**

Automated high-content imaging pipeline. PreciScan (module in Harmony™)-enabled low-magnification screening identifies condensate-positive cells for automated high-resolution confocal imaging and downstream segmentation, rapid condensate detection and quantitative analysis.

Representative KNIME workflow for high-content condensate metadata analysis and quantification. Workflow illustrating data ingestion, pre-processing, integration and downstream analysis of Operetta high-content imaging outputs. See Methods for details; full-version of KNIME workflows, including the details of the hidden tabs, are available from the authors upon reasonable request.

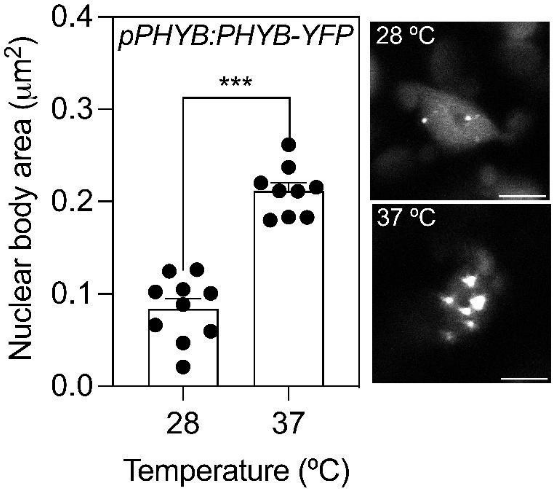


**Figure S5. Increased phyB NB size at hot temperatures in cotyledon cells.**

Confocal analysis of the area of the NBs in cotyledon cells driven by *pPHYB:PHYB-YFP*.

The seedlings were grown at 22°C, transferred to the indicated temperatures 1 h after the beginning of the photoperiod of the fourth day, and harvested 4 h after the beginning of temperature treatments. Scale bars: 5 µm. Means ±SE and individual values of 9-10 seedlings. Asterisks denote statistical significance of Student's *t* tests: ***, *P* <0.001.

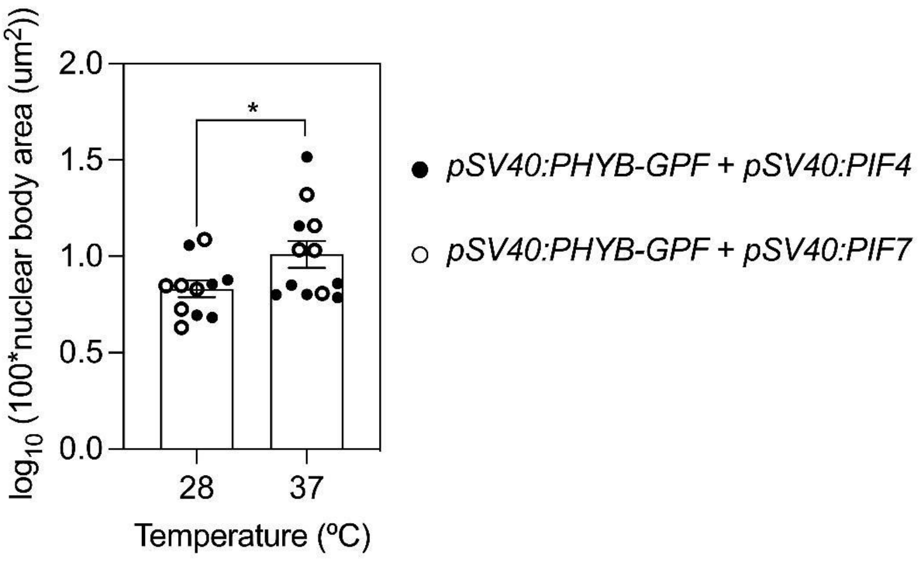


**Figure S6. High temperatures enhance phyB condensate size in mammalian cells.**

Confocal analysis of the area of the NBs driven by *pSV40:PHYB-eGFP* in CHO-K1 cells co-transfected with either *pSV40:PIF4-eCerulean* or *pSV40:PIF7-eCerulean.*

CHO-K1 cells were exposed to 15-min R followed by 60 min of darkness at the indicated temperatures. Data are the mean ±SE and individual values of 11-12 biological replicates. Data were log transformed to improve variance homogeneity. Numbers were multiplied by 100 before log transformation to improve the presentation (positive values); Y= 1 indicates 0.1 $\mu m^2$. Asterisks denote statistical significance of Student's *t* tests: *, $P < 0.05$.
