## Supplementary material for "Thermosensory pathway makes a U-turn to respond to heat": Dataset 1

**Quantification and normalisation of protein blots.** Data corresponding to the intensity of each band and its background and calculation of normalised PIF4-HA, PIF7-HA (4 h and 8 h), PHYB-GFP and PHYB-YFP abundance shown in Figure 4 and 5.

| | | PIF4-HA intensity | | Rubisco | | PIF4-HA/Rubisco | Blot average | Normalised PIF4-HA abundance |
|---|---|---|---|---|---|---|---|---|
| ºC | Replicate | Band | Background | Band | Background average | | | |
| 10 | 1 | 91.2 | 62.5 | 131.0 | 69.6 | 27.3 | 67.1 | 0.4 |
| 15 | 1 | 127.3 | 62.0 | 130.1 | 69.6 | 62.7 | 67.1 | 0.9 |
| 22 | 1 | 103.4 | 60.8 | 130.6 | 69.6 | 40.6 | 67.1 | 0.6 |
| 28 | 1 | 197.6 | 59.8 | 131.7 | 69.6 | 129.0 | 67.1 | 1.9 |
| 33 | 1 | 136.8 | 58.4 | 127.5 | 69.6 | 78.9 | 67.1 | 1.2 |
| 37 | 1 | 106.5 | 56.7 | 115.9 | 69.6 | 62.6 | 67.1 | 0.9 |
| 10 | 2 | 66.6 | 64.2 | 141.0 | 65.6 | 2.4 | 60.8 | 0.0 |
| 15 | 2 | 128.0 | 62.8 | 147.0 | 65.6 | 61.7 | 60.8 | 1.0 |
| 22 | 2 | 81.2 | 62.7 | 139.8 | 65.6 | 19.3 | 60.8 | 0.3 |
| 28 | 2 | 186.4 | 62.9 | 144.7 | 65.6 | 120.1 | 60.8 | 2.0 |
| 33 | 2 | 205.3 | 64.0 | 146.6 | 65.6 | 134.3 | 60.8 | 2.2 |
| 37 | 2 | 74.8 | 61.2 | 136.2 | 65.6 | 14.7 | 60.8 | 0.2 |

| 4 h | PIF7-HA intensity | | | | Rubisco | | PIF7-HA/Rubisco | | Blot average | Normalised PIF7-HA abundance | | |
|---|---|---|---|---|---|---|---|---|---|---|---|---|
| ºC | Replicate | Upper band | Lower band | Background | Band | Background average | Upper band | Lower band | | Upper band | Lower band | Sum of bands |
| 10 | 1 | 105.2 | 104.3 | 109.0 | 161.0 | 75.9 | -3.8 | -4.7 | 29.1 | -0.1 | -0.2 | -0.3 |
| 15 | 1 | 96.6 | 101.6 | 99.5 | 160.8 | 75.9 | -2.9 | 2.1 | 29.1 | -0.1 | 0.1 | 0.0 |
| 22 | 1 | 86.0 | 134.4 | 88.3 | 158.1 | 75.9 | -2.3 | 47.3 | 29.1 | -0.1 | 1.6 | 1.5 |
| 28 | 1 | 76.7 | 112.4 | 76.5 | 163.4 | 75.9 | 0.2 | 34.3 | 29.1 | 0.0 | 1.2 | 1.2 |
| 33 | 1 | 96.6 | 141.8 | 65.3 | 159.4 | 75.9 | 31.5 | 77.2 | 29.1 | 1.1 | 2.7 | 3.7 |
| 37 | 1 | 103.4 | 177.9 | 55.4 | 157.3 | 75.9 | 49.8 | 127.0 | 29.1 | 1.7 | 4.4 | 6.1 |
| 10 | 2 | 45.3 | 52.2 | 45.4 | 161.0 | 71.1 | -0.1 | 7.2 | 28.2 | 0.0 | 0.3 | 0.3 |
| 15 | 2 | 45.9 | 59.2 | 44.0 | 160.8 | 71.1 | 1.9 | 14.9 | 28.2 | 0.1 | 0.5 | 0.6 |
| 22 | 2 | 51.3 | 59.3 | 47.3 | 158.1 | 71.1 | 3.8 | 11.6 | 28.2 | 0.1 | 0.4 | 0.5 |
| 28 | 2 | 62.4 | 92.9 | 55.3 | 163.4 | 71.1 | 7.5 | 40.0 | 28.2 | 0.3 | 1.4 | 1.7 |
| 33 | 2 | 122.1 | 155.8 | 69.0 | 159.4 | 71.1 | 52.6 | 86.0 | 28.2 | 1.9 | 3.1 | 4.9 |
| 37 | 2 | 112.0 | 192.0 | 95.3 | 157.3 | 71.1 | 16.0 | 92.3 | 28.2 | 0.6 | 3.3 | 3.8 |
| 10 | 3 | 81.6 | 86.0 | 80.7 | 113.2 | 50.4 | 1.1 | 6.8 | 49.1 | 0.0 | 0.1 | 0.2 |
| 15 | 3 | 69.6 | 118.1 | 66.4 | 127.6 | 50.4 | 3.4 | 54.2 | 49.1 | 0.1 | 1.1 | 1.2 |
| 22 | 3 | 67.9 | 59.5 | 56.7 | 140.4 | 50.4 | 10.0 | 2.4 | 49.1 | 0.2 | 0.0 | 0.3 |
| 28 | 3 | 87.9 | 50.3 | 48.1 | 137.5 | 50.4 | 37.0 | 2.1 | 49.1 | 0.8 | 0.0 | 0.8 |
| 33 | 3 | 173.8 | 135.5 | 43.6 | 137.2 | 50.4 | 121.2 | 85.6 | 49.1 | 2.5 | 1.7 | 4.2 |
| 37 | 3 | 204.5 | 131.3 | 43.2 | 131.6 | 50.4 | 160.7 | 87.8 | 49.1 | 3.3 | 1.8 | 5.1 |
| 10 | 4 | 51.6 | 55.7 | 48.4 | 134.7 | 50.4 | 3.3 | 7.6 | 36.8 | 0.1 | 0.2 | 0.3 |
| 15 | 4 | 64.6 | 155.9 | 58.5 | 142.7 | 50.4 | 5.8 | 92.5 | 36.8 | 0.2 | 2.5 | 2.7 |
| 22 | 4 | 75.9 | 81.6 | 73.2 | 139.4 | 50.4 | 2.6 | 8.2 | 36.8 | 0.1 | 0.2 | 0.3 |
| 28 | 4 | 132.8 | 111.6 | 84.6 | 132.4 | 50.4 | 51.5 | 28.8 | 36.8 | 1.4 | 0.8 | 2.2 |
| 33 | 4 | 175.1 | 171.7 | 96.7 | 139.2 | 50.4 | 77.4 | 74.0 | 36.8 | 2.1 | 2.0 | 4.1 |
| 37 | 4 | 119.0 | 196.5 | 113.8 | 139.8 | 50.4 | 5.2 | 81.2 | 36.8 | 0.1 | 2.2 | 2.3 |

| 8 h | PIF7-HA intensity | | | | Rubisco | | PIF7-HA/Rubisco | | Blot average | Normalised PIF7-HA abundance | | |
|---|---|---|---|---|---|---|---|---|---|---|---|---|
| ºC | Replicate | Upper band | Lower band | Backgro und | Band | Backgrou nd average | Upper band | Lower band | | Upper band | Lower band | Sum of bands |
| 10 | 1 | 69.2 | 63.9 | 58.2 | 76.0 | 20.1 | 8.4 | 4.3 | 69.0 | 0.1 | 0.1 | 0.2 |
| 15 | 1 | 79.8 | 68.2 | 62.1 | 73.0 | 20.1 | 14.3 | 5.0 | 69.0 | 0.2 | 0.1 | 0.3 |
| 22 | 1 | 126.9 | 113.5 | 66.2 | 59.2 | 20.1 | 66.4 | 51.7 | 69.0 | 1.0 | 0.7 | 1.7 |
| 28 | 1 | 154.9 | 150.3 | 69.9 | 58.6 | 20.1 | 94.3 | 89.3 | 69.0 | 1.4 | 1.3 | 2.7 |
| 33 | 1 | 179.5 | 226.7 | 76.1 | 54.6 | 20.1 | 128.1 | 186.6 | 69.0 | 1.9 | 2.7 | 4.6 |
| 37 | 1 | 201.1 | 221.5 | 81.4 | 55.7 | 20.1 | 143.7 | 168.2 | 69.0 | 2.1 | 2.4 | 4.5 |
| 10 | 2 | 106.1 | 104.2 | 94.4 | 60.0 | 20.1 | 13.2 | 11.1 | 31.2 | 0.4 | 0.4 | 0.8 |
| 15 | 2 | 106.3 | 107.4 | 100.0 | 64.0 | 20.1 | 6.5 | 7.6 | 31.2 | 0.2 | 0.2 | 0.5 |
| 22 | 2 | 129.9 | 133.7 | 103.7 | 59.1 | 20.1 | 30.3 | 34.6 | 31.2 | 1.0 | 1.1 | 2.1 |
| 28 | 2 | 125.9 | 132.5 | 106.9 | 62.8 | 20.1 | 20.1 | 27.1 | 31.2 | 0.6 | 0.9 | 1.5 |
| 33 | 2 | 146.5 | 166.7 | 108.8 | 69.9 | 20.1 | 34.2 | 52.5 | 31.2 | 1.1 | 1.7 | 2.8 |
| 37 | 2 | 159.8 | 204.7 | 111.1 | 75.9 | 20.1 | 39.5 | 75.8 | 31.2 | 1.3 | 2.4 | 3.7 |
| 10 | 3 | 111.6 | 108.5 | 94.9 | 83.2 | 28.6 | 15.3 | 12.5 | 56.5 | 0.3 | 0.2 | 0.5 |
| 15 | 3 | 99.1 | 101.8 | 93.1 | 83.0 | 28.6 | 5.6 | 8.0 | 56.5 | 0.1 | 0.1 | 0.2 |
| 22 | 3 | 97.0 | 99.5 | 91.7 | 74.2 | 28.6 | 5.9 | 8.6 | 56.5 | 0.1 | 0.2 | 0.3 |
| 28 | 3 | 164.8 | 160.0 | 91.5 | 70.0 | 28.6 | 88.5 | 82.7 | 56.5 | 1.6 | 1.5 | 3.0 |
| 33 | 3 | 219.7 | 228.0 | 115.7 | 84.8 | 28.6 | 92.3 | 99.7 | 56.5 | 1.6 | 1.8 | 3.4 |
| 37 | 3 | 229.8 | 218.9 | 93.4 | 75.9 | 28.6 | 143.8 | 132.3 | 56.5 | 2.5 | 2.3 | 4.9 |
| 10 | 4 | 98.8 | 100.0 | 96.0 | 80.9 | 28.6 | 2.5 | 3.5 | 72.4 | 0.0 | 0.0 | 0.1 |
| 15 | 4 | 102.7 | 102.4 | 96.5 | 74.4 | 28.6 | 6.2 | 5.8 | 72.4 | 0.1 | 0.1 | 0.2 |
| 22 | 4 | 178.5 | 152.6 | 103.9 | 75.0 | 28.6 | 73.1 | 47.7 | 72.4 | 1.0 | 0.7 | 1.7 |
| 28 | 4 | 196.5 | 179.3 | 99.5 | 65.7 | 28.6 | 118.9 | 97.7 | 72.4 | 1.6 | 1.3 | 3.0 |
| 33 | 4 | 246.6 | 233.9 | 106.8 | 78.2 | 28.6 | 128.0 | 116.4 | 72.4 | 1.8 | 1.6 | 3.4 |
| 37 | 4 | 240.9 | 248.8 | 103.4 | 70.2 | 28.6 | 150.3 | 159.1 | 72.4 | 2.1 | 2.2 | 4.3 |

| | | phyB-GFP intensity | | | Rubisco | | phyB-GFP/Rubisco | | Blot average | Normalised phyB-GFP abundance | | |
|---|---|---|---|---|---|---|---|---|---|---|---|---|
| ºC | Replicate | Upper band | Lower band | Background | Band | Background average | Upper band | Lower band | | Upper band | Lower band | Sum of bands |
| 10 | 1 | 131.2 | 128.9 | 121.4 | 126.6 | 64.5 | 10.5 | 8.0 | 50.2 | 0.2 | 0.2 | 0.4 |
| 15 | 1 | 128.6 | 124.1 | 114.0 | 129.4 | 64.5 | 14.8 | 10.2 | 50.2 | 0.3 | 0.2 | 0.5 |
| 22 | 1 | 151.4 | 209.0 | 108.9 | 129.9 | 64.5 | 42.8 | 100.8 | 50.2 | 0.9 | 2.0 | 2.9 |
| 28 | 1 | 115.9 | 132.4 | 101.6 | 131.6 | 64.5 | 14.0 | 30.2 | 50.2 | 0.3 | 0.6 | 0.9 |
| 33 | 1 | 182.8 | 181.6 | 99.8 | 132.5 | 64.5 | 80.5 | 79.3 | 50.2 | 1.6 | 1.6 | 3.2 |
| 37 | 1 | 186.5 | 235.7 | 106.9 | 132.3 | 64.5 | 77.3 | 125.2 | 50.2 | 1.5 | 2.5 | 4.0 |
| 10 | 2 | 114.0 | 137.1 | 87.5 | 128.0 | 60.8 | 25.1 | 46.9 | 77.1 | 0.3 | 0.6 | 0.9 |
| 15 | 2 | 133.8 | 164.6 | 84.2 | 131.8 | 60.8 | 44.4 | 72.0 | 77.1 | 0.6 | 0.9 | 1.5 |
| 22 | 2 | 150.0 | 199.7 | 84.5 | 128.8 | 60.8 | 61.2 | 107.6 | 77.1 | 0.8 | 1.4 | 2.2 |
| 28 | 2 | 153.6 | 180.4 | 92.6 | 123.0 | 60.8 | 62.3 | 89.6 | 77.1 | 0.8 | 1.2 | 2.0 |
| 33 | 2 | 202.1 | 174.4 | 96.4 | 121.1 | 60.8 | 111.3 | 82.1 | 77.1 | 1.4 | 1.1 | 2.5 |
| 37 | 2 | 190.0 | 225.9 | 105.3 | 113.1 | 60.8 | 102.9 | 146.5 | 77.1 | 1.3 | 1.9 | 3.2 |
| 10 | 3 | 96.7 | 97.7 | 95.1 | 103.4 | 55.9 | 1.7 | 3.0 | 16.8 | 0.1 | 0.2 | 0.3 |
| 15 | 3 | 102.3 | 115.3 | 93.2 | 110.6 | 55.9 | 8.9 | 21.6 | 16.8 | 0.5 | 1.3 | 1.8 |
| 22 | 3 | 98.9 | 95.4 | 94.5 | 111.7 | 55.9 | 4.2 | 0.8 | 16.8 | 0.2 | 0.0 | 0.3 |
| 28 | 3 | 125.6 | 142.6 | 98.9 | 113.2 | 55.9 | 24.9 | 40.8 | 16.8 | 1.5 | 2.4 | 3.9 |
| 33 | 3 | 164.8 | 109.1 | 105.1 | 111.1 | 55.9 | 57.8 | 3.8 | 16.8 | 3.4 | 0.2 | 3.7 |
| 37 | 3 | 136.6 | 115.0 | 112.2 | 106.2 | 55.9 | 25.9 | 3.0 | 16.8 | 1.5 | 0.2 | 1.7 |
| 10 | 4 | 153.8 | 153.2 | 143.1 | 100.4 | 55.9 | 11.7 | 11.0 | 16.7 | 0.7 | 0.7 | 1.4 |
| 15 | 4 | 139.7 | 136.2 | 137.0 | 112.4 | 55.9 | 2.3 | -0.7 | 16.7 | 0.1 | 0.0 | 0.1 |
| 22 | 4 | 126.6 | 125.5 | 126.4 | 114.1 | 55.9 | 0.2 | -0.7 | 16.7 | 0.0 | 0.0 | 0.0 |
| 28 | 4 | 136.0 | 202.9 | 119.3 | 104.7 | 55.9 | 16.6 | 83.5 | 16.7 | 1.0 | 5.0 | 6.0 |
| 33 | 4 | 121.5 | 111.6 | 112.1 | 102.9 | 55.9 | 9.7 | -0.5 | 16.7 | 0.6 | 0.0 | 0.6 |
| 37 | 4 | 143.9 | 138.8 | 106.5 | 93.1 | 55.9 | 49.0 | 42.3 | 16.7 | 2.9 | 2.5 | 5.5 |

| | | phyB-YFP intensity | | | Rubisco | | phyB-YFP/Rubisco | | Blot average | Normalised phyB-YFP abundance | | |
|---|---|---|---|---|---|---|---|---|---|---|---|---|
| ºC | Replicate | Upper band | Lower band | Background | Band | Background average | Upper band | Lower band | | Upper band | Lower band | Sum of bands |
| 10 | 1 | 186.9 | 173.0 | 147.8 | 109.8 | 73.7 | 49.8 | 32.1 | 46.0 | 1.1 | 0.7 | 1.8 |
| 15 | 1 | 177.6 | 168.9 | 143.5 | 122.3 | 73.7 | 32.2 | 23.9 | 46.0 | 0.7 | 0.5 | 1.2 |
| 22 | 1 | 227.4 | 202.4 | 140.5 | 126.2 | 73.7 | 76.1 | 54.2 | 46.0 | 1.7 | 1.2 | 2.8 |
| 28 | 1 | 170.8 | 145.5 | 132.2 | 121.0 | 73.7 | 37.5 | 12.9 | 46.0 | 0.8 | 0.3 | 1.1 |
| 33 | 1 | 129.7 | 130.5 | 126.8 | 117.1 | 73.7 | 3.1 | 3.9 | 46.0 | 0.1 | 0.1 | 0.2 |
| 37 | 1 | 236.1 | 229.1 | 122.4 | 121.3 | 73.7 | 109.6 | 102.9 | 46.0 | 2.4 | 2.2 | 4.6 |
| 10 | 2 | 140.7 | 172.9 | 176.4 | 124.2 | 73.7 | 28.9 | 2.9 | 57.3 | 0.5 | 0.1 | 0.6 |
| 15 | 2 | 172.0 | 172.9 | 176.6 | 113.4 | 73.7 | 4.8 | 3.8 | 57.3 | 0.1 | 0.1 | 0.1 |
| 22 | 2 | 83.8 | 149.9 | 172.8 | 117.8 | 73.7 | 82.3 | 21.1 | 57.3 | 1.4 | 0.4 | 1.8 |
| 28 | 2 | 84.0 | 123.2 | 165.3 | 114.9 | 73.7 | 80.5 | 41.7 | 57.3 | 1.4 | 0.7 | 2.1 |
| 33 | 2 | 55.4 | 88.3 | 160.1 | 113.9 | 73.7 | 106.1 | 72.7 | 57.3 | 1.9 | 1.3 | 3.1 |
| 37 | 2 | 22.7 | 41.7 | 146.0 | 102.5 | 73.7 | 174.2 | 147.3 | 57.3 | 3.0 | 2.6 | 5.6 |
| 10 | 3 | 73.4 | 69.8 | 65.1 | 87.2 | 62.1 | 11.8 | 6.6 | 49.0 | 0.2 | 0.1 | 0.4 |
| 15 | 3 | 111.4 | 86.5 | 68.1 | 97.8 | 62.1 | 43.3 | 18.4 | 49.0 | 0.9 | 0.4 | 1.3 |
| 22 | 3 | 84.6 | 88.1 | 74.2 | 95.4 | 62.1 | 11.0 | 14.8 | 49.0 | 0.2 | 0.3 | 0.5 |
| 28 | 3 | 144.4 | 143.0 | 82.8 | 99.8 | 62.1 | 58.3 | 57.0 | 49.0 | 1.2 | 1.2 | 2.4 |
| 33 | 3 | 179.7 | 165.0 | 91.6 | 103.8 | 62.1 | 75.4 | 62.9 | 49.0 | 1.5 | 1.3 | 2.8 |
| 37 | 3 | 208.9 | 192.3 | 97.8 | 102.8 | 62.1 | 97.5 | 83.0 | 49.0 | 2.0 | 1.7 | 3.7 |
| 10 | 4 | 95.8 | 87.7 | 78.3 | 83.9 | 54.9 | 23.2 | 12.4 | 55.2 | 0.4 | 0.2 | 0.6 |
| 15 | 4 | 112.3 | 85.8 | 72.3 | 90.7 | 54.9 | 42.7 | 14.4 | 55.2 | 0.8 | 0.3 | 1.0 |
| 22 | 4 | 122.3 | 102.2 | 68.4 | 98.4 | 54.9 | 47.4 | 29.7 | 55.2 | 0.9 | 0.5 | 1.4 |
| 28 | 4 | 133.1 | 105.3 | 65.9 | 106.1 | 54.9 | 50.2 | 29.4 | 55.2 | 0.9 | 0.5 | 1.4 |
| 33 | 4 | 154.8 | 113.2 | 68.2 | 89.1 | 54.9 | 96.7 | 50.2 | 55.2 | 1.8 | 0.9 | 2.7 |
| 37 | 4 | 235.5 | 170.6 | 75.0 | 90.5 | 54.9 | 172.2 | 102.6 | 55.2 | 3.1 | 1.9 | 5.0 |
